# Early emergence for thermal processing in the neonatal somatosensory cortex and its perturbation in ASD

**DOI:** 10.1101/2025.10.09.681402

**Authors:** Ugo Zayan, Anastasia Ludwig, Nicolas Catz, Arthur Godino, Franck Jandard, Françoise Muscatelli, Claudio Rivera, Valery Matarazzo

## Abstract

Thermal perception is essential for neonatal adaptation and survival, yet the cortical processing during early development remains unknown. Here we show that non-painful cool, but not warm, orofacial stimulation evokes robust and lateralized activation in the whisker-related somatosensory cortex (wS1) of neonatal mice during the first postnatal week. These responses, revealed by mesoscopic in vivo imaging and electrophysiological recordings, partly overlap with tactile representations from the same region, indicating early regionalization of thermotactile maps. In a mouse model of autism spectrum disorder deficient for *Magel2*, we found altered spatial encoding of cool stimuli accompanied by increased local field potential activity, suggestive of cortical hyperexcitability. Our findings establish that the cortical processing of cool sensory information emerges from the first week of life and is disrupted in a genetic model of autism, highlighting early vulnerability of thermosensory circuits and their potential contributions to neurodevelopmental disorders.

## INTRODUCTION

Thermal perception is a fundamental aspect of sensory processing that enables organisms to adapt to fluctuating environmental conditions. Studies in adult rodents have elucidated a sophisticated network of temperature-sensitive receptors and brain regions ^1–13^, revealing that the somatosensory cortex (S1) and posterior insular cortex (pIC) are central to thermal integration ^1,2,8^.

These cortical regions support both the detection of subtle temperature variations and the integration of thermal with tactile information, making them critical for adaptive behavior ^14^. In particular, S1 is essential for cold perception and may facilitate thermotactile coding through shared representations with tactile input ^6,8^, while pIC has been implicated in broader temperature identification ^8^.

Despite significant advances in understanding thermoception in adult models, less is known about how these systems develop and function in early post-natal life.

This gap in knowledge is notable, as neonatal mice are highly dependent on thermal cues for survival, yet are unable to regulate their own core temperature.^15^.

During this period, sensory experience drives the maturation of cortical circuits and shapes functional architecture ^16–20^, as well as cognitive behavior ^21^, with whisker-mediated perception serving as a primary and major sensory system before the onset of vision and hearing ^22,23^. The whisker region of S1 (wS1, barrel cortex) emerges as a structurally and functionally defined area early in postnatal development, providing an ideal model for investigating the maturation of sensory processing ^24,25^.

Atypical sensory processing is recognized as an early and pervasive feature of autism spectrum disorders (ASD) ^26–29^ and is predictive of later social and cognitive difficulties ^30–33^ as well as the ASD phenotype severity ^34^. However, the developmental emergence of thermosensory processing—and its susceptibility to genetic perturbations relevant to ASD—remains unexplored. MAGEL2 gene is responsible for Schaaf-Yang syndrome, a complex neurodevelopmental characterized by ASD (ref: Tim Schubert and C. Schaaf, 2024))^35,36^. Previously, we showed that Magel2 knock-out (KO) mouse model displays early-life behavioral alterations in response to cool thermal stimuli^37^.

Here, we investigate for the first time the cortical integration of thermal information in neonatal mice, focusing on the maturation of wS1 networks. In parallel, we examine the presence of atypical thermosensory processing in the Magel2 mouse model of autism, which displays early-life behavioral alterations in response to cool thermal stimuli ^37^. This work elucidates the developmental trajectory of cortical thermosensation and its disruption in a genetic context relevant to neurodevelopmental disorders.

## RESULTS

### Mesoscale Neonatal Somatosensory Activation Following Tactile Stimulation in wil-type neonates

To examine mesoscale activity in the somatosensory cortex in pups (P6-P8, P: post-natal day) we used in-vivo 1-photon wide-field calcium imaging focusing on the wS1 region, a principal and functionally active area of this cortex during the first week of life ^24,25^. We performed manual whisker stimulation on awaked neonatal mice (P6-P8) expressing the calcium indicator GCaMP6f in cortical excitatory neurons (Fig.1a). Tactile stimulation consistently evoked robust GCaMP6f fluorescence responses temporally aligned with whisker deflections in awake P6–P8 animals (Fig. 1b), with latency and amplitude time-locked to stimulus onset in most trials (Fig. 1c). Neonates exhibited a high response rate of 90% to tactile stimulation (Fig. 1d). Quantitative analysis revealed an increase in cortical activity during stimulation relative to both baseline and post-stimulus intervals, as assessed by area under the curve per second (AUCs−1−1) at the animal (Fig. 1e) and trial levels (Extended Data Fig. 1a). Signal duration above threshold and peak amplitude during evoked responses were significantly elevated compared to non-stimulation periods across animals and trials (Extended Data Fig. 1b–e).

**Figure 1:**
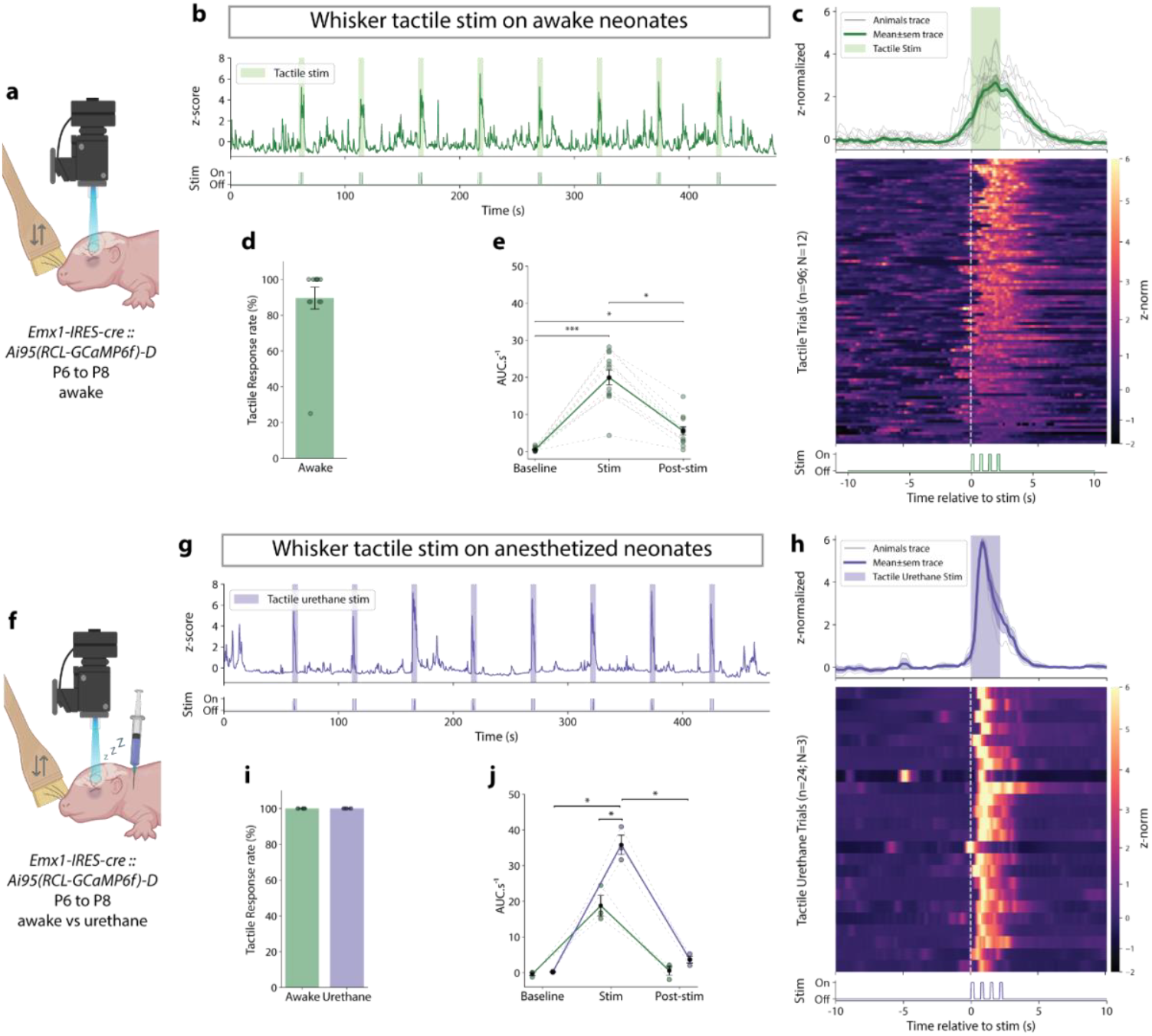
Mesoscale tactile evoked calcium events in S1 somatosensory cortex of neonates. **a**, Experimental schematic of mesoscopic1-photon calcium imaging in S1 somatosensory cortex during whisker tactile stimulation in awaked Emx1-IRES-cre::Ai95(RCL-GCaMP6f)-D neonates mice above (from P6 to P8; N=12 WT). **b**, Representative tactile evoked calcium events in the somatosensory cortex of one awake neonate receiving a series of 8 contralateral whisker tactile stimulations (green boxes). **c**, Top: Peristimulus time histogram (PSTH) of the evoked responses to tactile stimulations represented as the mean of all (green) or each (grey) animal. Middle: Heatmap of GCaMP6f signal of all tactile stimulations (trials) from all animals (n=96 trials for N=12 WT), aligned with the stimulation pulse (white dashed line). Bottom: representation of the manual back-and-forth whiskers stimulations. **d**, Animal average tactile evoked calcium response rate (N=12 WT). **e**, Quantification (Average area under the curve per second; AUC.s^-1^) for the three periods of recording: baseline, stim, post-stim. Friedman test p<0.0001 followed by Dunn post-hoc with Bonferroni adjustment: Baseline vs Stim p<0.0001, Baseline vs Post-stim p=0.0168, Stim vs Post-stim p=0.0283 (N=12 WT). **f** to **h**, follow the same description as **a** to **e** for the comparison of the GCaMP6f signal from mouse pups receiving tactile stimulation in an awake or anesthetized (urethane) state. **f**, Experimental schematic for urethane condition (N=3 WT for both conditions). **G**, Representative GCaMP6f signal from an anesthetized neonate recording. **h**, PSTH curve, and heatmap for all anesthetized animals (n=24 trials for N=3 WT). **i**, the Animal average tactile evoked calcium response rate in awake (green) or urethane (purple) condition (N=3 WT for both conditions). **j**, AUC.s^-1^ quantification for each period and conditions: awake (green) and urethane (purple). Two-way RM ANOVA followed by Bonferroni’s multiple comparisons test. Period effect p=0.0131; condition effect p=0.0194; interaction period x condition p=0.0009. For urethane condition: Baseline vs Stim p=0.0174, Stim vs Post-stim p=0. 0182. For the Stim period: Awake vs urethane p=0.0220 (N=3 WT for each condition). Data represented as mean ± sem. Significativity : * p < 0.05; ** p < 0.01; *** p < 0.001; n.s Non-significant.

We further assessed potential influences of age and sex on tactile-evoked activity. At all ages studied, robust evoked activity was still present; however, the overall amplitude of the calcium signal decreased as postnatal age increased, with significantly lower responses detected at postnatal day 8 compared to day 6, consistent with prior findings^38^ (Extended Data Fig. 1f–j). No significant differences between sexes were detected in the magnitude or pattern of the measured responses (Extended Data Fig. 1k–l).

To ensure that tactile-evoked cortical activity was not confounded by animal movement during stimulation, we repeated the experiments under urethane anesthesia (Fig. 1f). Anesthetized animals displayed enhanced and more widespread GCaMP6f responses during stimulation, with a concomitant reduction in spontaneous activity (Fig. 1g, h; Extended Data Fig. 2a, b). The response rate remained similar between awake and anesthetized groups (Fig. 1i), while the amplitude of the evoked signal during stimulation was significantly higher under urethane, both at the animal and trial levels (Fig. 1j; Extended Data Fig. 2c). Measures of response rate and magnitude above threshold were also significantly greater during stimulation than at baseline and post-stimulation, with improved response reliability under anesthesia (Extended Data Fig. 2d–g).

Collectively, these results demonstrate that robust, stimulus-locked whisker-evoked activity can be detected in wS1 during the first postnatal week and that anesthesia enhances the reliability and amplitude of these responses. These findings further validate the accuracy of our imaging region for subsequent analyses of thermally evoked activity in wS1 during early development.

### Mesoscale Neonatal Somatosensory Activation Following Cool, but Not Warm, Stimulation in wil-type neonates

After confirming our recording site in wS1, we performed cutaneous cooling of the snout (33°C to 15°C) in the same mice previously subjected to tactile stimulation (Fig. 2a). Cool stimulation reliably elicited increased GCaMP6f fluorescence in wS1, though with lower amplitude than responses evoked by whisker deflection (Fig. 2b). This cortical activation was tightly time-locked to the onset of cooling and exhibited a broader temporal profile compared to tactile responses (Fig. 2c). The response rate to cooling was approximately 55% (Fig. 2d), with significant increases in area under the curve per second (AUCs−1−1) during stimulation, observed in analyses pooled by animal and by trial (Fig. 2e, Extended Data Fig. 3a). Signal duration above threshold and peak response magnitude were both greater during the stimulus period, supporting the conclusion that peripheral cool stimulation can reliably drive activity in wS1 (Extended Data Fig. 3b–e).

**Fig 2:**
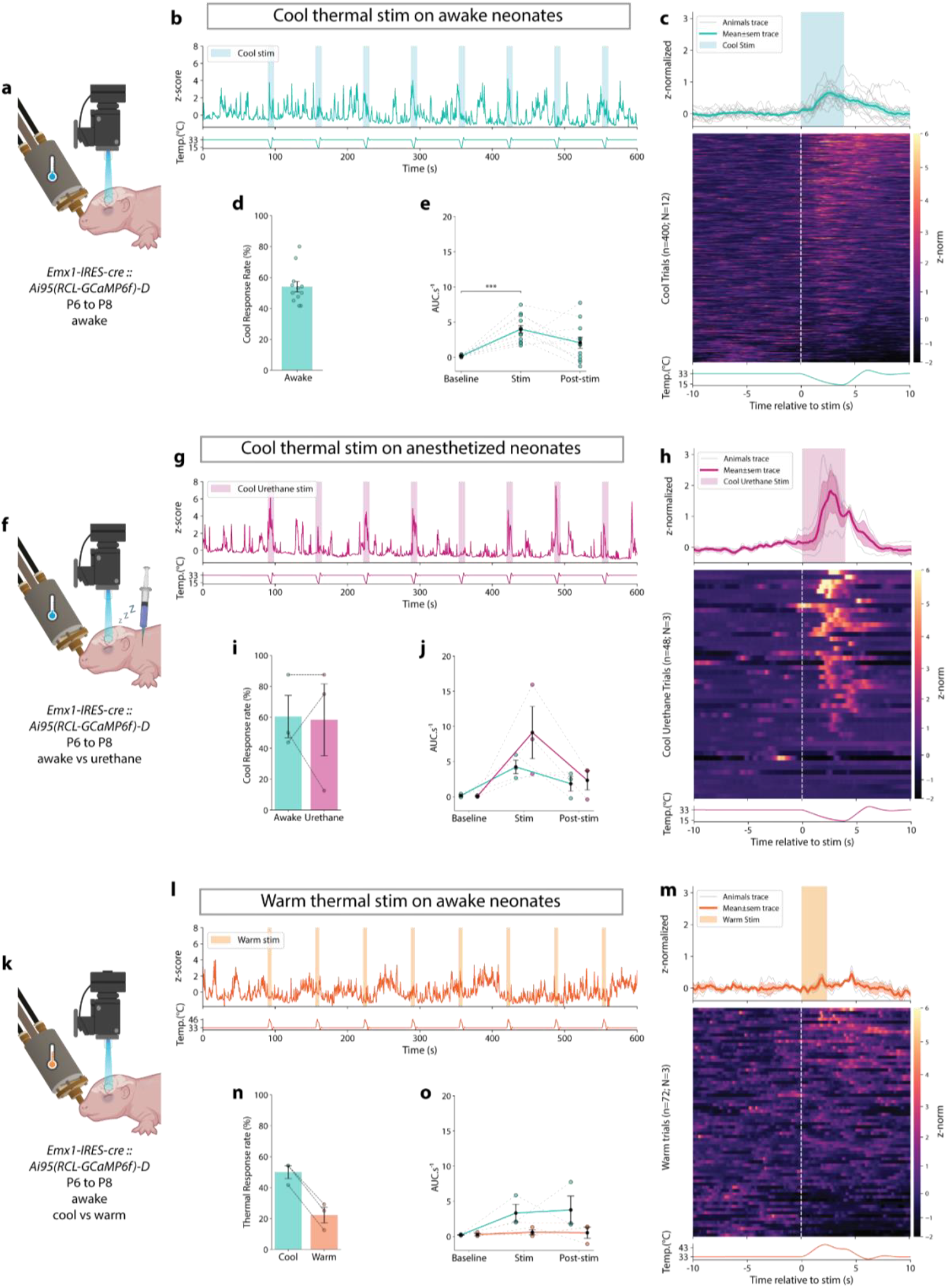
Mesocale thermal evoked calcium events in S1 somatosensory cortex of neonates. **a**, Experimental schematic of mesoscopic1-photon calcium imaging in S1 somatosensory cortex during topical orofacial cool stimulations in awaked *Emx1-IRES-cre::Ai95(RCL-GCaMP6f)-D neonates* (from P6 to P8; N=12 WT). **b**, Representative cool evoked calcium events in the somatosensory cortex of one awake neonate receiving a series of 8 contralateral cool stimulations (blue boxes). **c**, Top: PSTH of the evoked responses to cool stimulations represented as the mean of all (blue) and each (grey) animal receiving cool stimulation (blue box). Middle: Heatmap of GCaMP6f signal of all cool stimulations (all trials) from all animals (n=400 trials for N=12 WT), aligned with the stimulation pulse (white dashed line). Bottom: Orofacial skin temperature variations during cool stimulation. **d**, Animal average cool response rate l (N=12 WT). **e**, Quantification (AUC.s^-1^) of each cool stimulation of the three periods of recording: baseline, stim, and post-stim. Friedman tests *p=0*.*0023*, followed by post-hoc Dunn with Bonferroni adjustment: Baseline *vs* Stim *p=0*.*0002* (N=12 WT). **f** to **h** follows the same description as **a** to **e** for the comparison of the GCaMP6f signal of mouse pups receiving cool stimulation in an awake or anesthetized (urethane) state. **f**, Experimental schematic for urethane condition (N=3 WT for both conditions). **g**, Representative GCaMP6f signal from an anesthetized neonate recording. **h**, PSTH curve and heatmap of all trials from all anesthetized animals (n=48 trials for N=3 WT). **i**, the Animal average cool response rate for awake (blue) or urethane (magenta) condition. Wilcoxon test: Awake *vs* urethane *p=0*.*6547* (N=3 WT for both conditions). **j**, AUC.s^-1^ quantification for each period (baseline, Stim, Post-stim) and conditions: awake (blue) and urethane (magenta). Two-way RM ANOVA followed by Bonferroni’s multiple comparisons test. Period effect *p=0*.*1317*; condition effect *p=0*.*3605*; interaction period x condition *p=0*.*2865*. N=3 WT for both conditions. **k** to **o** follow the same description as **a** to **e** for the comparison of the GCaMP6f signal of mouse pups receiving cool and warm stimulations. **k**, Experimental schematic of warm stimulation (N=3 WT for both conditions). **l**, Representative GCaMP6f signal of one awake neonate recording during warm stimulations. **m**, PSTH curve and heatmap of all trials from all animals receiving warm stimulation (n=72 trials in N=3 WT). **n**, Animal average thermal response rate after cool (blue) and warm (orange) stimulations. Wilcoxon test: Cool *vs* Warm *p=0*.*2500* (N=3 WT for both conditions). **o**, AUC.s^-1^ quantification for each period (baseline, Stim, Post-stim) and conditions: cool (blue) and warm (orange). Two-way RM ANOVA followed by Bonferroni’s multiple comparisons test. Period effect *p=0*.*4035*; condition effect *p=0*.*0935*; interaction period x condition *p=0*.*2519* (N=3 WT for both conditions). Data represented as mean ± sem. Significativity : * p < 0.05; ** p < 0.01; *** p < 0.001; n.s Non-significant.

Interestingly, unlike tactile stimulation, neither the magnitude nor response rate of cool-evoked activity varied significantly with age, and no sex differences were observed (Extended Data Fig. 3f–o).

To rule out movement artefacts as a source of cool-evoked responses, we conducted identical experiments under urethane anesthesia (Fig. 2f). The magnitude of cortical activity evoked by cooling was even greater under anesthesia, with responses roughly twice as large and displaying sharper kinetics (Fig. 2g, h; Extended Data Fig. 4a, b). Although anesthesia did not significantly alter the probability of cool-responsive trials (Fig. 2i; Extended Data Fig. 4c), it significantly elevated AUCs−1−1 and response magnitude for both animals and trials (Fig. 2j; Extended Data Fig. 4d–h), while preserving the specificity of increased activity during stimulation versus baseline.

Given evidence in adult mice that cooling, but not warming^8^, evokes robust S1 responses, we next tested whether thermal sensitivity is similarly tuned in neonates. Warm stimulation (33°C to 46°C) applied to the same skin region did not produce time-locked increases in GCaMP6f signal (Fig. 2l, m; Extended Data Fig. 4i, j). Although animal-based response rates for cool and warm conditions were not significantly different (Fig. 2n), trial-based response analysis revealed a significant difference, with only cool stimulation producing robust increases in AUCs−1−1 and response parameters (Fig. 2o; Extended Data Fig. 4k–p).

To compare somatotopic organization, we assessed receptive field overlap for tactile and thermal stimulation of the same region. Stereotactic mapping verified recording consistency (Extended Data Fig. 5a), with thermal stimulation activating broader receptive fields than tactile, but with ∼30% spatial overlap between modalities (Extended Data Fig. 5b, c).

Together, these results establish that neonatal mice exhibit robust, reproducible wS1 activation in response to cool, but not warm, cutaneous stimulation. This cool-evoked activity is amplified by anesthesia, confirming it is not artefactual, and overlaps partially with tactile-driven cortical fields. Our findings suggest that, as in adults, cold is a salient modality driving neonatal somatosensory cortical activity and reveal the functional substrates for early thermo-tactile integration in wS1.

### Network-Level Characterization of Tactile-Evoked Activity in wS1 cortex of wil-type neonates

Having established robust calcium responses to tactile and cool stimulation in neonatal wS1, we next sought to delineate the underlying neural network dynamics. To this end, we performed in vivo extracellular recordings with silicon probes targeted to layer 4 (L4) of S1 in awake wild-type pups (P6–P8), delivering precisely controlled whisker stimulations using a piezoelectric actuator (Fig. 3a). Analysis of multiunit activity (MUA) and local field potential (LFP) revealed clear tactile-evoked responses, demonstrated by increased neuronal spiking and LFP amplitude across specific channels following stimulation (Fig. 3b). Raster plots from representative recordings demonstrated a marked, phase-locked increase in spike events aligned to stimulus onset (Fig. 3c).

**Fig 3:**
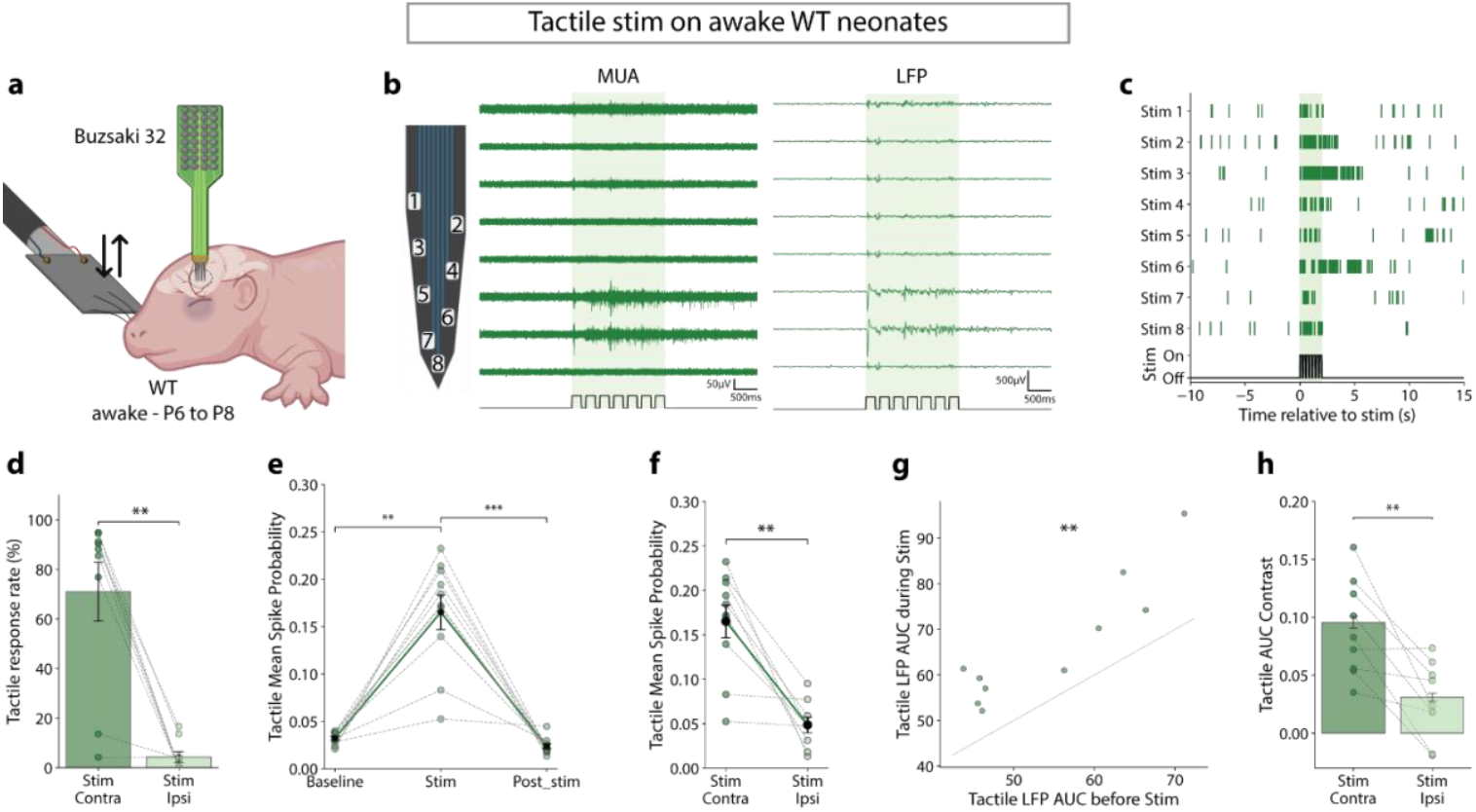
Cortical tactile evoked activity in S1 of WT neonates. **a**, Experimental schematic of electrophysiological extracellular recordings in S1 somatosensory cortex during whisker tactile stimulation in awake WT mice from P6 to P8; N=10 WT. **b**, Schematic of one of the 4 shanks of the silicon probes. Representative tactile MUA and LFP evoked activity in S1 for a tactile stimulation (green box) performed on an awake WT newborn. Traces from one shank are organized from the shallowest to the deepest channel. **c**, Raster plot of an example cluster for 8 tactile stimuli. The stimulation is represented by the green box. **d**, Mean tactile response rate quantification for the contra- and the ipsilateral stimulation. Wilcoxon test *p=0*.*0078* (N=9 WT). **e**, Mean tactile spike probability quantification for the three recording periods. Friedman test *p=0*.*0004* followed by Dunn post-hoc with Bonferroni adjustment: Baseline *vs* Stim *p=0*.*0075*, Stim *vs* Post-stim *p<0*.*0001* (N=10 WT). **f**, Mean tactile spike probability quantification for the contra- and the ipsilateral stimulation. Wilcoxon test *p=0*.*0039* (N=9). **g**, Quantification of the tactile AUC of the derived LFP signal for the Baseline and Stim periods. Each point represents one animal. Wilcoxon test *p=0*.*002* (N=10 WT). **h**, Quantification of the tactile AUC-derived LFP signal contrast (Stim to Baseline signal) for the contra- or ipsilateral side. Each point represents an animal. Wilcoxon test *p=0*.*0078* (N=9 WT). Data represented as mean ± sem. Significativity : * p < 0.05; ** p < 0.01; *** p < 0.001; n.s Non-significant.

To verify that these responses were not driven by movement artefacts, we compared stimulation of contralateral and ipsilateral whiskers in awake animals. Quantification of response rates indicated significantly stronger cortical activation following contralateral whisker stimulation (Fig. 3d). Assessment of tactile-evoked spiking using spike density functions showed significant increases in probability during stimulation compared to both baseline and post-stimulus periods (Fig. 3e), with contralateral stimulation producing the most robust spike probability (Fig. 3f).

Further, analysis of the LFP revealed that area under the curve (AUC) values were significantly elevated following whisker stimulation for contralateral input, both at the animal and channel levels (Fig. 3g; Extended Data Fig. 6a). Calculation of the post-stimulus AUC contrast reinforced the lateralized nature of the response, with greater evoked activity for the contralateral side (Fig. 3h; Extended Data Fig. 6b).

Together, these data corroborate our mesoscale calcium imaging results and extend previous studies by showing that neonatal cortical activity in response to whisker stimulation is already present at the single-unit and population levels^39–42^, displaying clear lateralization for the contralateral side. These findings also confirm precise probe localization within wS1.

### Neonatal Somatosensory Cortex Networks encode Cool but Not Warm Stimuli with Lateralized Activation in wild-type neonates

We next assessed whether wS1, which encodes tactile information at early postnatal stages, similarly processes thermal input. Simultaneous extracellular recordings in the same region as previous tactile experiments revealed that cool (but not warm) thermal stimulation of the snout reliably evoked multiunit (MUA) and local field potential (LFP) activity (Fig. 4a–e). Although minor numerical differences were evident between cool and warm stimuli or between contra- and ipsilateral stimulation (Fig. 4f), cool, but not warm, stimulation elicited a significant increase in spike probability for contralateral input, paralleling tactile stimulation patterns (Fig. 4g). Cool stimulation produced a significant rise in spike probability during though this effect was weaker than that seen with tactile input, consistent with the lower response rate for cool (3.5%) compared to tactile stimulation (70%) (Fig. 4h). Warm stimulation did not induce any MUA activity (Fig. 4i).

**Fig 4:**
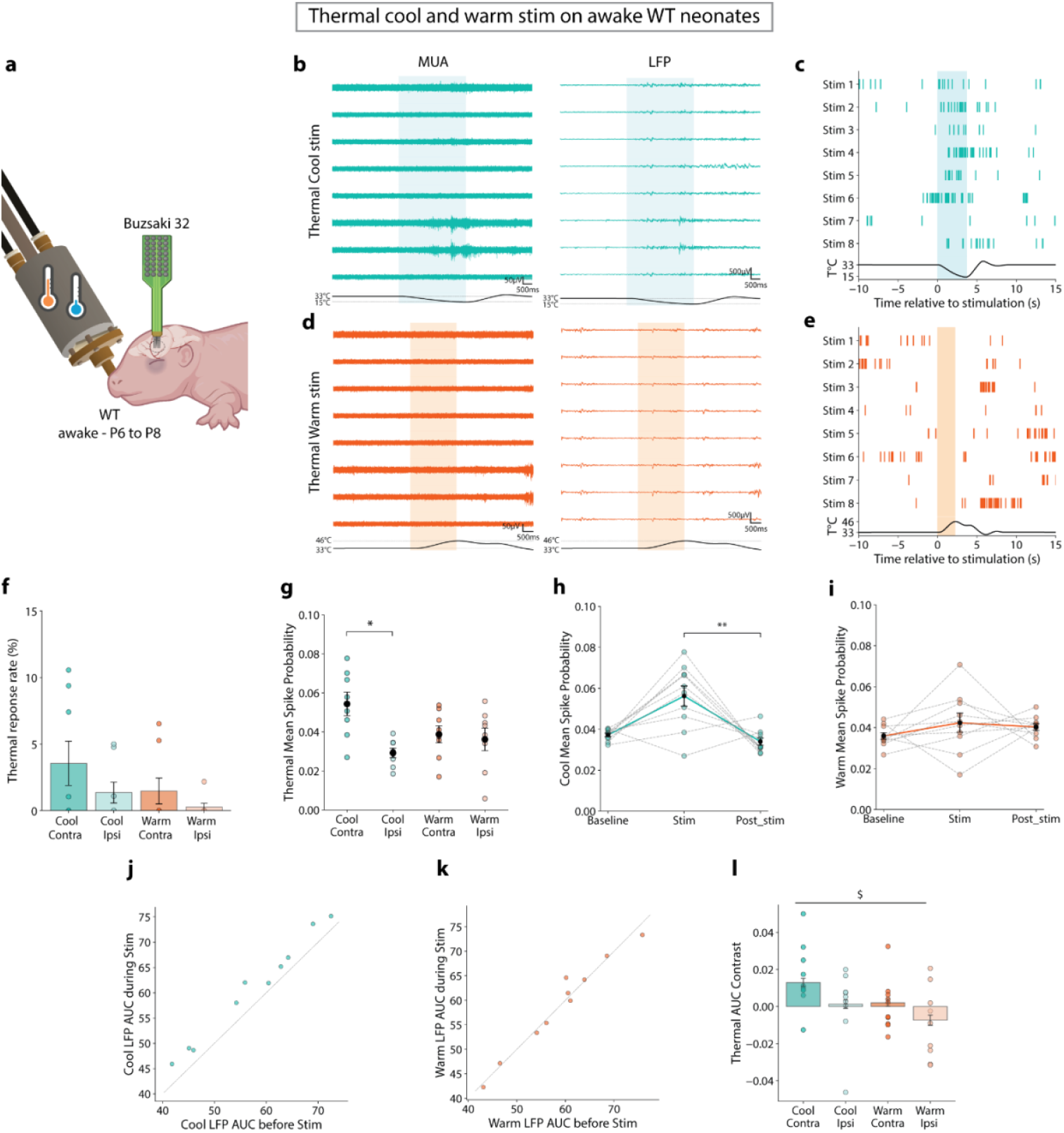
Thermal cool cortical evoked activity in S1 of WT neonates. **a**, Experimental schematic of electrophysiological extracellular recordings in S1 cortex during topical orofacial cool or warm stimulation in awake WT mice from P6 to P8; N=10 WT. **b**, Representative thermal evoked MUA and LFP signal for a cool stimulation (blue box). Traces from one shank are organized from the shallowest to the deepest channel. **c**, Raster plot of an example cluster with 8 cool stimuli (blue box). **d**, same as **a** for the thermic warm stimulation (orange box). **e**, same as **c** for the thermic warm stimulation (orange box). **f**, Mean thermal response rate quantification for cool, warm, contralateral, and ipsilateral stimulations. Two-way RM ANOVA followed by Bonferroni’s multiple comparisons test. Temperature effect *p=0*.*2310*; laterality effect *p=0*.*0556*; interaction temperature x laterality *p=0*.*4640*. (N=8 WT for all conditions). **g**, Mean thermal spike probability quantification for cool, warm, contralateral, and ipsilateral stimulations. Two-way RM ANOVA followed by Bonferroni’s multiple comparisons test. Temperature effect *p=0*.*4157*; laterality effect *p=0*.*0394*; interaction temperature x laterality *p=0. 0349*. Cool-contra *vs* Cool-ipsi *p=0*.*0409*. (N=8 WT for all conditions). **h**, Mean thermal cool spike probability quantification for the three periods of recording. Friedman test *p=0*.*0074* followed by Dunn post-hoc with Bonferroni adjustment: Stim *vs* Post-stim *p=0*.*0015* (N=10 WT). **i**, Same as **h** for warm stimulations. Friedman test *p=0*.*2725*. (N=10 WT). **j**, Cool AUC-derived LFP signal quantification for the Baseline and Stim periods. Each point represents one animal. Wilcoxon test *p=0*.*002* (N=10 WT). **k**, same as **j** for warm stimulation. Wilcoxon test *p=0*.*6953* (N=10 WT). **l**, Quantification of the thermal AUC-derived LFP signal contrast (Stim to Baseline) for the cool, warm contra- and ipsilateral stimulation. Each point represents an animal. Mixed-effects analysis followed by Bonferroni’s multiple comparisons test. Temperature effect *p=0*.*0674*; laterality effect *p=0*.*0459*; interaction temperature x laterality *p=0*.*6216*. (N=10 WT). The main effect of the lateralization is noted as $. Data represented as mean ± sem. Significativity : * p < 0.05; ** p < 0.01; *** p < 0.001; n.s Non-significant.

Analysis of LFP contrast (AUC difference pre- and post-stimulation) indicated no specific effects of stimulus temperature or side (Fig. 4j; Extended Data Fig. 6c). Nevertheless, AUC increased robustly following cool stimulation but not warm, confirming that only cool stimulation reliably activates wS1 (Fig. 4k–l; Extended Data Fig. 6d–e).

Together, these data demonstrate that, during early development, cool—but not warm—sensory input selectively activates wS1 local networks. Cool-evoked cortical responses are weaker than tactile responses, mirroring results observed in adult forepaw stimulation^6^, and an established lateralization at this stage. No evidence of warm-evoked activity was observed in either MUA or LFP, further supporting the specificity of neonatal wS1 for cool stimuli.

### Early Life Altered Thermosensory Network Responses in Magel2 KO neonates

Atypical sensory processing is a hallmark of autism spectrum disorder (ASD) ^29^, emerging early in development and commonly observed in affected individuals ^26–28^. Since our previous work demonstrated an altered neonatal behavioral responses to cool thermal stimulation in Magel2 KO pups, we further investigated whether these phenotypes extend to cortical network function. Thus, we assessed thermosensory coding in Magel2 KO pups using in vivo electrophysiological recordings (Fig. 5a).

**Fig 5:**
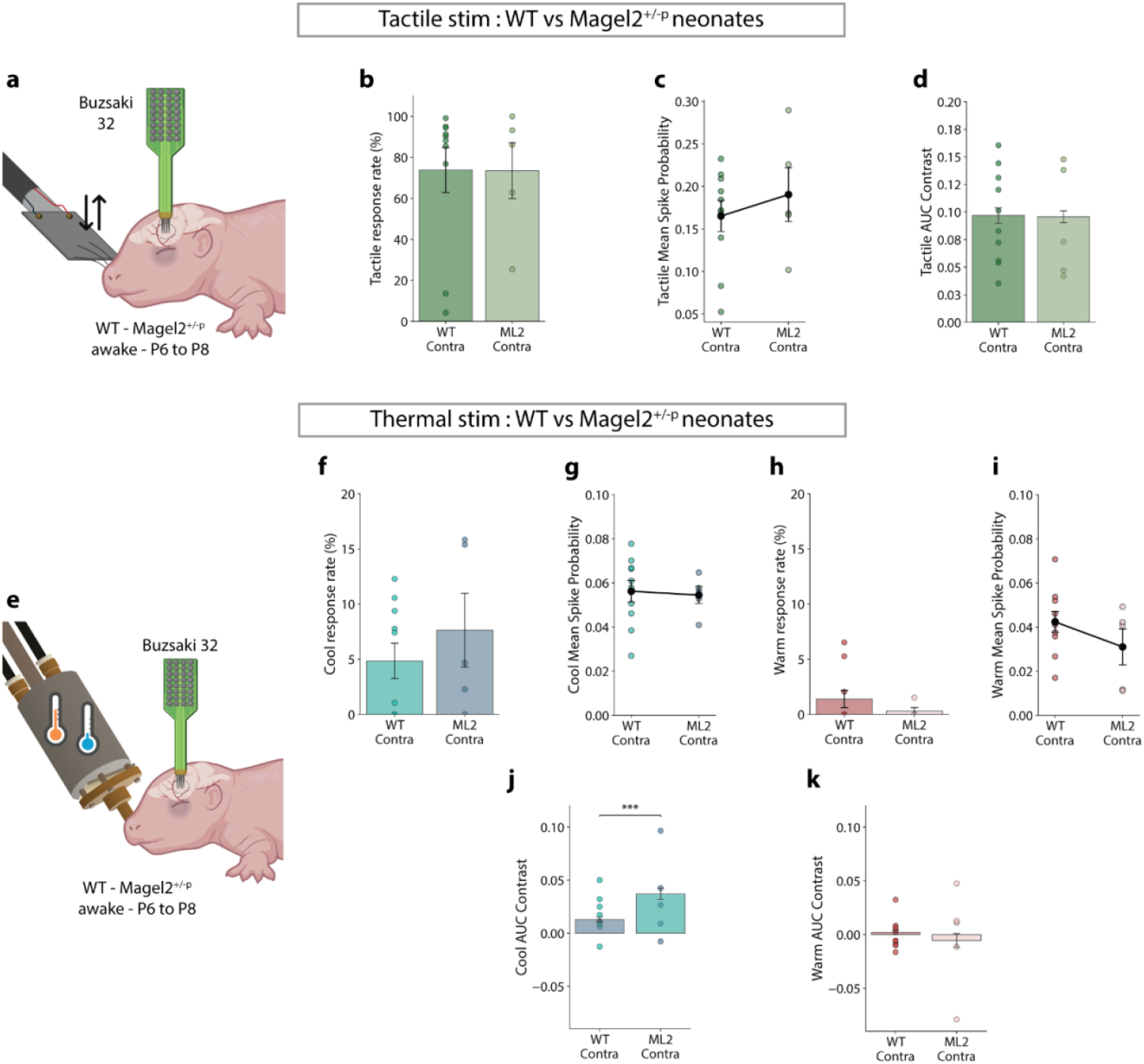
Tactile and thermal evoked activity comparison between WT and Magel2 KO neonates. **a**, Experimental schematic of electrophysiological extracellular recordings in S1 cortex during whisker tactile stimulation in awake WT or Magel2 KO mice from P6 to P8. N=10 WT and N=5 Magel2 KO. **b**, Mean tactile response rate quantification for WT and Magel2 KO neonates. Mann-Whitney test *p=1*.*0000* (N=10 WT and N=5 Magel2 KO). **c**, Mean tactile spike probability quantification for WT and Magel2 KO neonates. Mann-Whitney test *p=0*.*8591* (N=10 WT and N=5 Magel2 KO). **d**, Quantification of the tactile AUC-derived LFP signal contrast (Stim to Baseline) for WT and Magel2 KO neonates. Each point represents one animal. Mann-Whitney test *p=0*.*625* (N=10 WT, N=5 Magel2 KO). **e**, Experimental schematic for topical orofacial thermal cool or warm stimulation in awake WT or Magel2 KO mice from P6 to P8 (N=10 WT, N=5 Magel2 KO). **f**, Mean thermal cool response rate for WT and Magel2 KO neonates. Mann-Whitney test *p=0*.*4175* (N=10 WT and N=5 Magel2 KO). **g**, Mean thermal cool spike probability for WT and Magel2 KO neonates. Mann-Whitney test *p=0*.*5941* (N=10 WT and N=5 Magel2 KO). **h**, Same as **f** for warm stimulation. Mann-Whitney test *p=0*.*5822* (N=10 WT and N=5 Magel2 KO). **i**, Same as **g** for warm stimulation. Mann-Whitney test *p=0*.*3097* (N=10 WT and N=5 Magel2 KO). **j**, Quantification of the cool AUC-derived LFP signal contrast for WT and Magel2 KO neonates. Each point represents an animal. Mann-Whitney test *p<0*.*0001* (N=10 WT, N=5 Magel2 KO). **k**, same as **j** for thermal warm stimulation. Mann-Whitney test *p=0*.*4602* (N=10 WT, N=5 Magel2 KO). Data represented as mean ± sem. Significativity : * p < 0.05; ** p < 0.01; *** p < 0.001; n.s Non-significant.

Under whisker stimulation, Magel2 KO neonates displayed robust wS1 cortical activation, as indicated by increases in multiunit activity (MUA) and local field potential (LFP) signals, with response rates preferentially lateralized to the contralateral side (Extended Data Fig. 7a–c). MUA analysis revealed a trend toward higher spike probability during stimulation, again more pronounced for the contralateral input (Extended Data Fig. 7d,e). LFP analysis showed a significant increase in contrast (post-vs. pre-stimulation AUC) for contralateral versus ipsilateral whisker deflection at the animal and trial level (Extended Data Fig. 7f,g). LFP AUC values were consistently elevated following whisker deflection across animals and trials (Extended Data Fig. 7h,i), confirming wS1 probe placement and the presence of tactile-evoked responses in Magel2 KO neonates. Comparisons with wild-type (WT) littermates revealed no significant differences in MUA response rate, spike probability, or LFP AUC contrast following contralateral tactile stimulation (Fig. 5b–d).

Having established normal tactile-evoked wS1 activity in Magel2 KO pups, we tested thermal responsiveness by delivering identical cool and warm orofacial stimuli. Consistent with WT controls, Magel2 KO neonates showed wS1 network activation (increased MUA and LFP) in response to cool, but not warm, stimulation (Extended Data Fig. 8a–d). Thermal stimulation response rates and spike probabilities did not differ significantly between types or stimulation sides (Extended Data Fig. 8e). Notably, Magel2 KO pups lacked the laterality effect in spike probability seen in WT controls (Extended Data Fig. 8f), although both genotypes exhibited similar contralateral cool response rates and spike probabilities (Fig. 5f,g). The increase in spike probability between stimulation and post-stimulation periods observed with contralateral cool stimulation in WT pups was absent in Magel2 KO animals (Extended Data Fig. 8g). Warm stimulation failed to increase MUA activity or response rate in either genotype (Extended Data Fig. 8h; Fig. 5h,i).

LFP analyses of thermal-evoked responses in Magel2 KO neonates revealed no difference in AUC contrast for different types and sides of stimulation (Extended Data Fig. 8i). Cool, but not warm, stimulation produced AUC increases for the majority of animals, with weaker effects observed for warm stimuli (Extended Data Fig. 8j–n). Importantly, the AUC contrast following contralateral cool— but not warm—stimulation was significantly higher in Magel2 KO pups than in WT suggesting an increase of synaptic input activities (Fig. 5j,k).

Together, these findings reveal that Magel2 KO neonates exhibit intact tactile-evoked wS1 cortical responses, comparable to WT littermates, but display network-level alterations in the thermosensory domain. Specifically, Magel2 KO neonates show increased LFP responses and a deficit in spatially selective (lateralized) coding of cool stimuli. Those data suggest altered network function at the cortical wS1 cortical level for thermosensory processing.

## DISCUSSION

The development of sensory systems, including thermoception, is fundamental to an organism’s ability to interact with its environment. Investigating how thermal processing emerges during early life is crucial for understanding both typical sensory maturation and the influence of environmental or genetic perturbations on these pathways. Given the centrality of thermal cues for neonatal survival and adaptation, our study provides insights into the critical windows and vulnerabilities of early sensory development.

We demonstrate that the somatosensory cortex (wS1) of neonatal mice is already capable of robustly and selectively responding to non-noxious cool, but not warm, orofacial stimulation, with clear lateralization to the contralateral side. This functional specificity is present from the first postnatal week and is sustained under anesthesia, highlighting the early maturation of thermosensory cortical circuits. Consistent with work in adult rodents^6,8^, warm stimulation did not elicit significant wS1 activation, suggesting conserved modality selectivity across the lifespan. The very small amplitude GCaMP signal that we saw might be explained by a lick of signal coming from S2 known to respond to warm ^43^. The partial overlap between tactile- and cool-evoked cortical maps further implies that regional organization of sensory processing is established early in development.

Considering that atypical thermosensory sensitivity is observed in ASD patients ^44–47^, we examined a genetic relevant ASD model. We observed that Magel2 KO neonates showed no impairment in tactile-evoked cortical signaling but exhibited disruption of spatial coding for cool stimulation, reflected by loss of lateralized discrimination and increased local field potential (LFP) activity in wS1. This increase in LFP amplitude may indicate hyperexcitability or altered synaptic integration, a phenotype paralleled by the atypical thermosensory behaviors observed in this model ^37^. Notably, similar associations between cortical hyperexcitability, aberrant sensory processing, and behavioral hyporesponsiveness have previously been reported only in adult ASD mouse models ^48–51^. More generally, atypical cortical sensory activity has often been associated with sensory-altered behaviors in ASD models ^52–54,54,55^. Our findings extend these observations to the early postnatal period, suggesting that aberrant thermosensory network development is an early feature of the Magel2 KO phenotype.

Collectively, this work establishes that functional cool thermosensation is present in neonatal cortex and is susceptible to disruption in a model of ASD. Our results highlight the importance of early sensory circuit maturation and suggest that thermosensory network vulnerability may contribute to atypical processing and behavioral phenotypes in neurodevelopmental disorders. Future studies should address the longitudinal trajectory of these deficits and their molecular underpinnings, as well as the relevance of these findings to human sensory development and ASD.

## ACKNOWLEDGMENTS

The project leading to this publication has received funding from the Fondation Lejeune (VM 2022-00137); Foundation Recherche Medicale (UZ Phd), MERNT (UZ PhD), Mobility Grant from the GIS Autism.

This work received support from the French government under the France 2030 investment plan, as part of the Initiative d’Excellence d’Aix-Marseille Université - A*MIDEX (AMX-19-IET-007) through the Marseille Maladies Rares Institute (MarMaRa)

Thanks to Stefania Sarno from the Century Turing Centre for Living Systems (Aix Marseille Univ, CNRS, INSERM, Marseille, France) for her help regarding electrophysiological analyses. Some of the illustrations in figures have been created using Biorender (VM licence)

## METHODS

### Mice

Mice were handled and cared for following the Guide for the Care and Use of Laboratory Animals (N.R.C., 1996) and the European Communities Council Directive of September 22nd, 2010 (2010/63/EU,74). Experimental protocols were approved by the institutional Ethical Committee Guidelines for animal research with the APAFIS accreditation no. #34564-2022010612088993 from the French ministry of higher education, research and innovation. All efforts were made to minimize the number of animals used. For calcium imaging, C57BL/6J Emx1-IRES-Cre females have been crossed with Ai95(RCL-GCaMP6f)-D male to express the GCaMP6f in pyramidal neurons of the cortex. Mice from P6 to P8 have been recorded. For extracellular electrophysiological recordings, we recorded WT C57BL/6J mice (The Jackson Laboratory) and C57BL/6J Magel2 KO mice obtained by crossing WT females with Magel2^tm1.1Mus−/−^ males. Mice, from P6 to P16, were maintained on a 12:12 h light/dark inverted cycle (07:00 lights off; 19:00 lights on) and were provided with food and water ad libitum.

### Stereotaxic surgeries

For Calcium imaging recordings, 45 minutes before the surgery, mice were subcutaneously (S.C.) injected with Buprenorphine (0.03mg/kg), and lidocaine cream was applied to the head of the animal for analgesia. Pups were placed and head-fixed in a stereotaxic apparatus (Kopf Instruments). The head of the animal was cleaned and sterilized using a mix of Iodine and ethanol at 70%. Pups were anesthetized with isoflurane: 3.5% induction 5-10 min; 2% during skin removal and skull cleaning; 1% while mounting the holder; with 90% oxygen all along. 0.5mg/ml lidocaine + adrenaline were applied to the skin cuts for 1 min and then removed. The holder (see Suchkov et al. 2022) ^56^ was placed above the somatosensory cortex using both stereotaxic coordinates and targeting the medial cerebral arteria bifurcation. Then it was fixed to the skull using a bit of cyanoacrylate glue and then with dental cement. Pups remain in the stereotactic frame with airflow and 0% isoflurane until the cement solidifies and the pup starts moving. Then, pups are transferred into a heating chamber set at 35°C for recovery. During the totality of the surgery, mice were on a heating pad set at 37°C.

For the electrophysiological recordings, the same surgical steps were followed as for the calcium imaging recordings. However, the support system was different. It consisted of a plastic cylinder glued to the base of the animal’s skull with dental cement, into which two metal rods could be inserted to hold the head in place. In addition, a cylindrical dental cement stud was built around the recording area with two cannulas inside to allow constant perfusion of 37°C heated aCSF during the recording. A circular craniotomy of 1 mm diameter is made over the recording area using the tip of an insulin syringe. The bone is removed, and the dura is left intact for the recording. The brain is immersed and heated throughout the recording session.

In both protocols, the animals are allowed 45 minutes to 1 hour to recover from surgery, depending on the animal’s condition.

### Thermostimulation device (Thermod)

Two thermal heads were made. One with a Peltier module of 4×4 mm (cooling power = 0.8W) and the other 6×6 mm (cooling power = 1.2W). Both thermal heads are equipped with two temperature sensors. The main one, a Pt100/4-wire, is glued to the top of the Peltier module and is in direct contact with the mouse. The second, a 100kOhm NTC thermistor, is glued to the heatsink. The temperature difference is used to keep the Peltier module within a safe operating range. The Peltier module is powered by a Thermo Electric Cooling temperature controller from Meerstetter Engineering (TEC-1091). This controller has a true bipolar DC power source for cooling and heating up to ±4A at ±21Vdc. The supplied PC software allows Peltier configuration and temperature control via a USB connection. Standard fixed, periodic (sinus, square, sawtooth), and arbitrary curves are easily programmable. An additional custom-printed circuit board has been added to monitor the main temperature sensor in parallel with TEC 1091. This voltage is shifted and scaled to give a reading of 0V to 5V for a temperature of 0°C to 50°C. This signal is sent to the auxiliary analog input of the Neuronexus SmartBox and allows the temperature values to be synchronized with the data from the electrophysiological probes.

### Stimulation protocol

For the manual tactile stimulation paradigm, the protocol was a 60s baseline followed by 2s stimulation (4x up and down with a Q-tip). A recording consists of 8 stimulations with 50s rest between stimulations. For the automated tactile stimulation paradigm, a Pizzo system composed of Piezoceramic Plates (PI France) controlled by a Piezo Amplifier was used (E-650.00 LVPZT-AMPLIFIER, PI France). A piece of Q-tip was placed at the end of the Piezoceramic Plates, and an up-and-down motion was performed to stimulate the animal’s whiskers. The same timing was used, with 2s of stimulation corresponding to 7 stimulations of 200ms at 100ms intervals. A recording consists of 8 stimulations with 50s rest between stimulations. For the thermic cool stimulation paradigm, Thermode was set at 33°C and took 2s to go from room temperature to 33°C and stayed for 120s. Then the stimulation started and the thermode went to 15°C in 3.9s, stayed for 2s, and then went back to 33°C in 2s. A recording consists of 8 stimulations with 60s rest at 33°C between stimulations. For the thermic warm stimulation paradigm, Thermode was set at 33°C and took 2s to go from room temperature to 33°C and stayed for 120s. Then the stimulation started and the thermode went to 46°C in 2.1s, stayed for 2s, and then went back to 33°C in 2s. A recording consists of 8 stimulations with 60s rest at 33°C between stimulations.

### Drug treatment

For anesthesia during the recordings, we injected urethane at 1.3mg/g with 10µl/g split in 2 doses injected in s.c with 15 min between the 2 doses. At the end of calcium recordings, an i.p. injection of 50µl of solution with pentobarbital was used for euthanasia, and stereotaxic coordinates of the recording area were taken.

### In-vivo 1-photon wide-field calcium imaging recordings and analysis

Animals were placed on the recording setup inside a box to maintain as much darkness as possible during the recording. They were placed on a 37°C heating pad and covered with a cotton blanket. They were head-restrained using the system described by Suchkov et al., 2022, which allowed us to perfuse the top of the recording area with 37°C heated PBS and maintain a stable brain temperature. We performed the manual tactile or thermal stimulation protocol described above, with a 5-minute rest between recordings. Urethane injections were performed without moving the animal from the setup to maintain the same focus. To record the GCaMP6f signals from the somatosensory cortex, custom-made software (InVigilo Neurotar) was used. Images were taken with a custom-made Neurotar wide-field microscope: camera Andor Zyla 4.2 with an X4 objective (ThorLab TL4X 0.2 NA) and an exposure time of 90 ms for the FITC channel. The effective size of the window was (HxW): 512×512 with binning 4×4 and a resolution of 15.192 microns per pixel. The acquisition rate of the camera and the thermod software was 10 Hz. The acquisition rate of the Neuronexus SmartBox pro 1kHz and the To synchronize the recording with thermal stimulation, Master 8, triggered by the calcium imaging acquisition software, was used to trigger the thermode software and the Neuronexus SmartBox pro, which received a TTL signal from both the camera and thermode.

Post-acquisition analyses were performed using custom Python code (GitHub). First, we used the ImageJ software to find the receptive field of the somatosensory cortex that responded to our stimulation (tactile or thermal). We used a custom ImageJ macro to do this. Once the receptive field was found for each stimulation and recording, we extracted and converted the data into fluorescence time series using Python scripts (GitHub) and associated them with the time of stimulation (tactile stimulation) or temperature (thermal stimulation). Then, to compare activity between animals and recording sessions, time series data from individual animals were analyzed. The bleaching slope of each animal’s recordings was analyzed to check for any need for baseline correction. To analyze the GCaMP6f signals, we made a standardization using a z-score (zF = (F – mean(F)) / std(F)). To study the time-locked cerebral activity concerning the stimulation, a period of 10s before and after the onset of the stimulation was selected, and the normalized (z-norm) signal was calculated by subtracting the z-score of these selected periods from the Mean of the respective baseline z-score. Based on the temperature profile for thermal stimulation or the timing for manual tactile stimulation, we divided the selected period into 3 sub-periods. Baseline, which starts 10s before the start of stimulation, Stimulation, and Post-stimulation, which starts immediately after the end of stimulation and lasts until 10s after the start of stimulation. For peri-event analysis, signal trace data are quantified and averaged for each trial (stimulation) of one animal. Then the mean z-norm of all animals is calculated. For the Heatmap of trials, we calculated the sum of z-norm during the stimulation period for each trial, and we sorted them from the highest to the lowest z-norm sum values.

To calculate the response rate to the different stimulations, we first calculated the z-score distribution of all animals and recordings combined with the excluded stimulation period. We took the 95th percentile of this distribution as our threshold (1.86). To consider that an animal responded to stimulation, we checked that during the stimulation period at least one z-norm value was above the threshold. To complete this analysis, we calculated two further metrics: the above threshold rate and the response magnitude. The above threshold rate corresponds to the number of values above the threshold for each period (baseline, stim, and post-stim) divided by the total number of values for these periods. In other words, the time spent above the threshold for each period. Response magnitude is the sum of the z-norm values above the threshold divided by the number of values in each period. We calculated the area under the curve per second (AUC.s-1) for each period by subtracting all negative values from the positive ones. We then normalized this AUC by dividing it by the duration of each period. Quantification of response rate, AUC.s-1, above threshold rate, and response magnitude were calculated for each period of the different types of stimulation with N=animals and n=trials (stimulation). The same analysis was performed for animals receiving tactile/thermal cold stimulation in the awake or anesthetized (under urethane) state and for those receiving thermal cold and warm stimulation, making a paired comparison between conditions.

### Calcium imaging Statistics

The Wilcoxon test was used to compare data from 2 paired groups, such as tactile-awake and tactile-urethane, cool-awake and cool-urethane, or cool and warm. The Mann-Whitney test was used to compare data from 2 independent groups, such as sex. Friedman’s test followed by Dunn’s post-hoc test with Bonferroni correction was used to compare data from multiple paired groups, such as for periods. The Kruskal-Wallis test was used to compare multiple independent groups, such as age. The scipy.stats and scikit-post-hocs Python packages were used for these tests. Two-way repeated measures ANOVA followed by Bonferroni’s multiple comparisons test was used to compare the effect of two variables, such as period with age, sex, or condition for a n of animals. Those tests were performed on GraphPad Prism 8.0.2. For the same type of comparison with n trials, more complex multifactorial designs were used on R v4.2.2 to account for the nesting of the data. Linear mixed models were calculated using the R lmer Test function for mixed effects models, and random effects (conceptualizing nested observations at 3 levels, such as trial, record, and animal) were modeled as random. Subsequent analysis of variance (LMM-ANOVA) was performed using type III sums of squares with the Kenward-Roger approximation of degrees of freedom. Post-hoc tests were performed using the R emmeans package, and significance was adjusted using the Bonferroni method at the 0.05 level. For example, to analyze the effect of periods and age on the AUC.s-1, the model is: *AUC*.*s-1 ∼ Period * Age + (1* | *Trial) + (1* | *Rec) + (1* | *Animal)*. Bar and line graphs represent mean ± sem. Significance was set at p < 0.05 with * p < 0.05; ** p < 0.01; *** p < 0.001; n.s non-significant.

### In-vivo electrophysiological extracellular recordings and analysis

Animals were placed on the recording setup, on a 37°C heating pad, and covered with a cotton blanket. They were head-restrained using a custom-made system described in the surgical section. The brain was maintained at physiological temperature using a 37°C heated aCSF solution perfused into the dental cement chamber. Silicon probes are implanted into the wS1 cortex at 407µm depth to target neurons of layer 4. We performed automated tactile all-whiskers stimulation using the piezoelectric element, which allows us to perform stereotyped and time-locked stimulation as described in the stimulation protocol section. The thermal stimulation protocol was the same as that used for calcium imaging. At the end of the recordings, brains were collected for histological analysis to verify the implantation site of the DiI-labelled silicon probes. These recordings were made using the Neuronexus SmartBox system coupled with the acute SmartLink A32 headstage and Buzsaki32 silicon probes. For this system, the recordings are stored in.xdat format.

To analyze the spike signal, we performed spike sorting using Spyking Circus. To run our files through the sorter, we converted them into a.dat format using custom-written Python code. Once the sorting was done, we performed curation on each file. We first eliminated all clusters based on a list of criteria. All clusters with a maximum amplitude lower than 26µV (our value of baseline noise during the recordings), with a spike count between 30 and 2500, were associated with noise. We then made the classification of each remaining cluster based on the auto-correlogram (refractory period of 1.5 ms), the mean waveform of the spike, the inter-stim interval (ISI), the firing rate, and the position of the signal on the probe map. These selected clusters are then processed independently using custom Python scripts. First, the spike times for each stimulation are aligned with the stimulus onset as time 0 and a window of 10s before and after the stimulus to compare the temporal activity of the cluster. A convolution method was used to transform the spike time into a series of Gaussians, 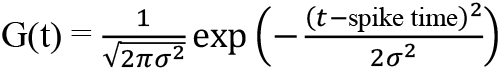, with a sigma of 100ms, which were summed to obtain a spike density function (SDF) matched to each stimulus for each cluster. The SDF is then normalized to give an area under the curve of 1 for each stimulation. This gave us a spike probability for each cluster that was averaged over the 8 stimuli. We then calculated the distribution of SDF values over the baseline and post-stim period for each cluster and determined a significance threshold based on the 95th percentile of this distribution. For each cluster, we checked whether there was a mean SDF value during the stim period that was above the threshold to consider that cluster to be responsive to stimulation. We were then able to calculate the response rate for each type of stimulation for each animal. We also calculated for each animal the mean SDF of each period (baseline, stim, and post-stim) of each stimulation from each cluster. Data were presented with an N=animal and n=clusters.

For LFP analysis, the signal for each channel of the electrode was smoothed with a Savinsky-Golay filter, derived, and rectified. The area under the curve (AUC) of these processed responses was used as a measure of the amount of activation. The data have been aligned with the stimulation, and the contrast of the AUC has been calculated by subtracting the AUC of one condition from another for each electrode. All analyses were carried out in the same way, regardless of the type of stimulation.

### In-vivo electrophysiological extracellular statistics

The Wilcoxon test was used to compare data from 2 paired groups for tactile, cool, and warm stimuli. The Mann-Whitney test was used to compare data from 2 independent groups. Friedman’s test followed by Dunn’s post-hoc test with Bonferroni correction was used to compare data from multiple paired groups, e.g., for periods. The Python packages scipy.stats and scikit_post-hocs were used for these tests. Two-way repeated measures ANOVA or Mixed model analysis, followed by Bonferroni’s multiple comparisons test were used to compare the effect of two variables, such as temperature and stimulation side. These tests were performed using GraphPad Prism 8.0.2. Bar and line graphs represent mean ± SEM. Significance was set at p < 0.05 with * p < 0.05; ** p < 0.01; *** p < 0.001; n.s non-significant.

## Data availability

All data, including raw calcium imaging and electrophysiological data, are available on request from the corresponding author. Both will be made public upon publication - for review, please email the corresponding author for private access keys.

## Code availability

The custom Python and R codes used in this study, including statistical analysis for both calcium imaging and electrophysiological data, are available upon request.

## EXTENDED DATA

**Extended Data Fig. 1:**
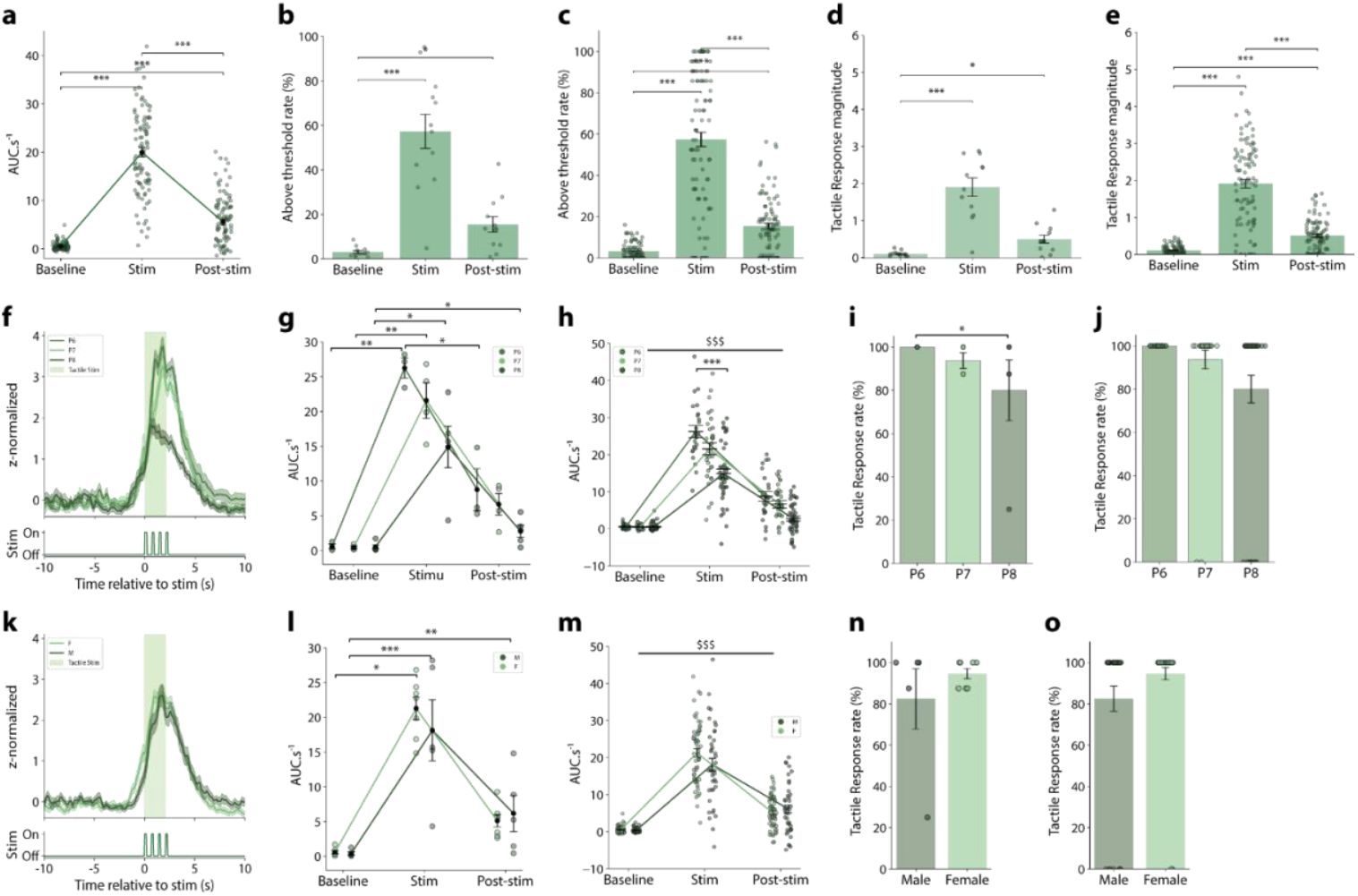
Sex and age effect on mesoscale tactile evoked calcium events in neonatal S1 cortex. **a**, Average area under the curve per second (AUC.s^-1^) quantification for each tactile stimulation period (Baseline, Stim, and Post-stim). Friedman test *p<0*.*0001*, followed by Dunn post-hoc with Bonferroni adjustment: Baseline *vs* Stim *p<0*.*0001*, Baseline *vs* Post-stim *p<0*.*0001*, Stim *vs* Post-stim *p<0*.*0001* (n=96 trials). **b**, Animal average above a threshold rate of the GCaMP6f signal for each tactile stimulation period. Friedman test *p=0*.*0001*, followed by Dunn post-hoc with Bonferroni adjustment: Baseline *vs* Stim *p<0*.*0001*, Baseline *vs* Post-stim *p=0*.*0238* (N=12 WT). **c**, All trials average above the threshold rate of the GCaMP6f signal for each tactile stimulation period. Friedman test *p<0*.*0001*, followed by Dunn post-hoc with Bonferroni adjustment: Baseline *vs* Stim *p<0*.*0001*, Baseline *vs* Post-stim *p<0*.*0001*, Stim *vs* Post-stim *p<0*.0001 (n=96 trials). **d**, Animal average tactile response magnitude of the GCaMP6f signal for each tactile stimulation period. Friedman test *p=0*.*0001*, followed by Dunn post-hoc with Bonferroni adjustment: Baseline *vs* Stim *p<0*.*0001*, Baseline *vs* Post-stim *p=0*.*02*, Stim *vs* Post-stim *p=0*.*0992* (N=12 WT). **e**, All trials average tactile response magnitude of the GCaMP6f signal for each tactile stimulation period. Friedman test *p<0*.*0001*, followed by Dunn post-hoc with Bonferroni adjustment: Baseline *vs* Stim *p<0*.*0001*, Baseline *vs* Post-stim *p<0*.*0001*, Stim *vs* Post-stim *p<0*.*0001* (n=96 trials). **f**, Top: PSTH curve of GCaMP6f signal during tactile stimulation (green box) for animals sorted by age: P6 (medium green), P7 (light green), and P8 (dark green). Bottom: representation of manual whiskers back-and-forth stimulations. **g**, Animal mean ± sem of AUC.s^-1^ quantification for each period and age. Two-way RM ANOVA followed by Bonferroni’s multiple comparisons test. Period effect *p<0*.*0001*; age effect *p=0*.*0249*; interaction period x age *p=0*.*0607*. For age P6: Baseline *vs* Stim *p=0*.*0098*. For age P7: Baseline *vs* Stim *p=0*.*0093*. For age P8: Baseline *vs* Stim *p=0*.*0267;* Stim *vs* Post-stim *p=0*.*0174*. For period Stimulation: P6 *vs* P8 *p=0*.*0484* (P6: N=3; P7: N=4; P8: N=5 WT). **h**, Trials mean ± sem of AUC.s^-1^ quantification for each period and age. LMM-ANOVA followed by Bonferroni-adjusted post-hoc tests. Period effect *p< 0*.*0001*; age effect *p=0*.*0249*; interaction period x age *p<0*.*0001*. For ages P6 and P7, every comparison of periods has *p<0*.*0001*. For age P8; Stim *vs* Baseline and Stim *vs* Post-stim have also *p<0*.*0001*. The main effect of the period is noted as $$$. For the period of stimulation; P6 *vs* P8 *p=0*.*0008* (P6: n=24; P7: n=32; P8: n=40 trials). **i**, Animal average tactile response rate sorted by age. Kruskal Wallis test: *p=0*.*2535* (P6: N=3; P7: N=4; P8: N=5 WT). **j**, Trials average tactile response rate sorted by age. Kruskal Wallis test: *p=0*.*0267*, followed by Dunn post-hoc with Bonferroni adjustment: P6 *vs* P8 *p=0*.*0350* (P6: n=24; P7: n=32; P8: n=40 trials). **k** to **o** follows the same description as **f** to **j** for the study of the sex effect on the GCaMP6f signal of mouse pups receiving tactile stimulation. **k**, Top: Peristimulus curve of GCaMP6f signal during tactile stimulation (green box) for all animals sorted by sex: Female (F) (light green) and Male (M) (dark green). **l**, AUC.s^-1^ quantification for each period and sex. Two-way RM ANOVA followed by Bonferroni’s multiple comparisons test. Period effect *p<0*.*0001*; sex effect *p=0*.*7163*; interaction period x age *p=0*.*4317*. For males: Baseline *vs* Stim *p=0*.*0446*. For females: Baseline *vs* Stim *p<0*.*0001*; Baseline *vs* Post-stim *p=0*.*0073* (females: N=7; males: N=5 WT). **m**, same as in **l** with n number of trials. LMM-ANOVA followed by Bonferroni-adjusted post-hoc tests. Period effect *p<0*.*0001*; sex effect *p=0*.*7163*; interaction period x sex *p=0*.*0329*. For females and males, every comparison of periods has *p<0*.*0001*. The main effect of the period is noted as $$$ (females: n=56; males: n=40 trials). **n**, Average tactile response rate for animals sorted by sex. Mann-Whitney test *p=0*.*9262* (females: N=7; males: N=5 WT). **o**, same as in **n** with n number of trials. Mann-Whitney test *p=0*.*0570* (females: n=56; males: n=40 trials). Data represented as mean ± sem. Significativity : * p < 0.05; ** p < 0.01; *** p < 0.001; n.s Non-significant.

**Extended Data Fig. 2:**
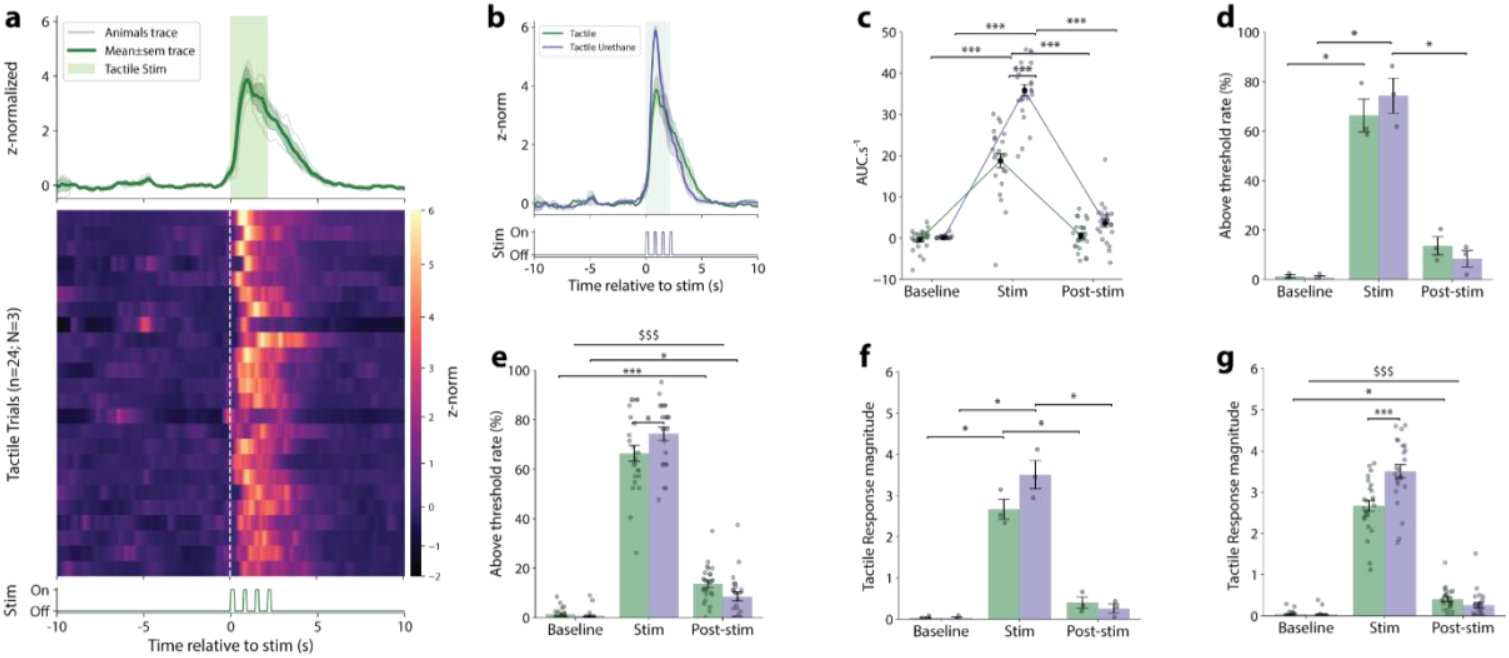
Mesocale tactile evoked activity in the S1. **A**, Top: Peristimulus curve of the mean ± sem GCaMP6f signal during tactile stimulation (green box) for all (green) and each (grey) animal. Middle: Heatmap of GCaMP6f signal for all tactile stimulation (trials) from all awake animals (n=24 trials in N=3 animals), aligned with the stimulation beginning (white dashed line). Bottom: representation of manual back-and-forth whisker stimulations. **B**, Top: Peristimulus curve of the mean ± sem GCaMP6f signal for animals under awake (green) or urethane (purple) condition during the tactile stimulation (green box). Bottom: representation of manual back-and-forth whisker stimulations. **C**, Average AUC.s^-1^ quantification for each period of the tactile stimulation and the two conditions: awake (green) and urethane (purple). LMM-ANOVA followed by Bonferroni-adjusted post-hoc tests. Period effect *p<0*.*0001*; condition effect *p<0*.*0001*; interaction period x condition *p<0*.*0001*. For conditions awake and urethane, Stim *vs* Baseline and Stim *vs* Post-stim have *p<0*.*0001*. For the period of stimulation, awake *vs* urethane *p<0*.0001 (n=24 trials for each condition). **D**, Average above threshold rate of the GCaMP6f signal for each period and condition: awake (green) and urethane (purple). Two-way RM ANOVA followed by Bonferroni’s multiple comparisons test. Period effect *p=0*.*0021*; condition effect *p=0*.*6341*; interaction period x condition *p=0*.*1353*. For awake condition: Baseline *vs* Stim *p=0*.*0310*. For urethane condition: Baseline *vs* Stim *p=0*.*0325*, Stim *vs* Post-stim *p=0. 0176* (N=3 WT for each condition). **E**, same as in **D** with n number of trials. LMM-ANOVA followed by Bonferroni-adjusted post-hoc tests. Period effect *p<0*.*0001*; condition effect *p=0*.*624*; interaction period x condition *p=0*.*002*. For condition awake and urethane, Stim *vs* Baseline and Stim *vs* Post-stim have *p<0*.*0001* noted as a main effect of the period with $$$. For the awake condition, Baseline *vs* Post-stim *p<0*.*0001*, and for the urethane condition, Baseline *vs* Post-stim *p=0*.*0341*. For the period of stimulation, awake *vs* urethane *p=0*.0241 (n=24 trials for each conditions). **F**, Average tactile response magnitude of the GCaMP6f signal for each period and condition: awake (green) and urethane (purple). Two-way RM ANOVA followed by Bonferroni’s multiple comparisons test. Period effect *p=0*.*0036*; condition effect *p=0*.*0322*; interaction period x condition *p=0*.*0081*. For awake condition: Baseline *vs* Stim *p=0*.*0279* and Stim *vs* Post-stim *p=0*.*0440*. For urethane condition: Baseline *vs* Stim *p=0*.*0326* and Stim *vs* Post-stim *p=0*.*0240* (N=3 animals for both conditions). **G**, same as in **F** with n number of trials. LMM-ANOVA followed by Bonferroni-adjusted post-hoc tests. Period effect *p<0*.*0001*; condition effect *p=0*.*0018*; interaction period x condition *p<0*.*0001*. For condition awake and urethane, Stim *vs* Baseline and Stim *vs* Post-stim have *p<0*.*0001* noted as a main effect of the period with $$$. For awake condition, Baseline *vs* Post-stim *p=0*.*0383*. For the period of stimulation, awake *vs* urethane *p<0*.*0001* (n=24 trials for each conditions). Data represented as mean ± sem. Significativity : * p < 0.05; ** p < 0.01; *** p < 0.001; n.s Non-significant.

**Extended Data Fig. 3:**
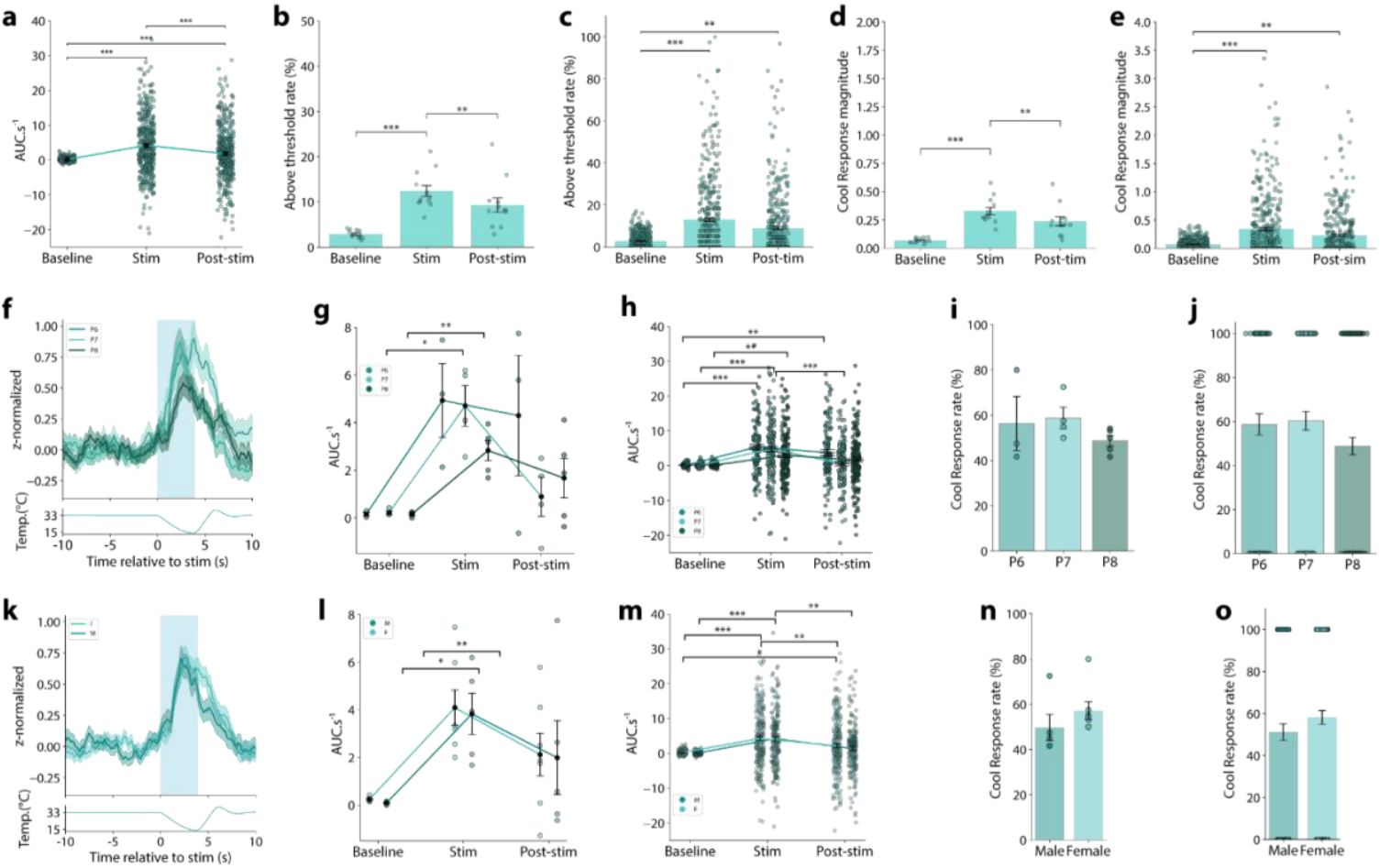
Sex and age effects on thermal cool evoked activity. **a**, AUC.s^-1^ quantification for each period of thermal cool stimulation. Friedman test *p<0*.*0001*, followed by Dunn post-hoc with Bonferroni adjustment: Baseline *vs* Stim *p<0*.*0001*, Baseline *vs* Post-stim *p<0*.*0001*, Stim *vs* Post-stim *p<0*.*0001* (n=400 trials). **b**, Average above threshold rate of the GCaMP6f signal for each period of thermal cool stimulation. Friedman test *p=0*.*0001*, followed by Dunn post-hoc with Bonferroni adjustment: Stim *vs* Baseline *p<0*.*0001*, Stim *vs* Post-stim *p=0*.*0036* (N=12 WT). **c**, same as in **b** with n number of trials. Friedman test *p<0*.*0001*, followed by Dunn post-hoc with Bonferroni adjustment: Baseline *vs* Stim *p<0*.*0001*, Baseline *vs* Post-stim *p<0*.*0001*, Stim *vs* Post-stim *p<0*.*0001* (n=96 trials). **d**, Average cool response magnitude of the GCaMP6f signal for each period of cool stimulations. Friedman test *p<0*.*0001*, followed by Dunn post-hoc with Bonferroni adjustment: Stim *vs* Baseline *p<0*.*0001*, Post-stim *vs* Baseline *p=0*.*0039* (n=12 WT). **e**, same as in **d** with n number of trials. Friedman test *p<0*.*0001*, followed by Dunn post-hoc with Bonferroni adjustment: Baseline *vs* Stim *p<0*.*0001*, Baseline *vs* Post-stim *p=0*.*0031* (n=400 trials). **f**, Top: Peristimulus curve of the mean ± sem GCaMP6f signal during thermal cool stimulation (blue box) for all animals sorted by age: P6 (dark turquoise), P7 (light turquoise), and P8 (deep turquoise). Bottom: Temperature variations during the cool stimulation. **g**, AUC.s^-1^ quantification for each period and age. Two-way RM ANOVA followed by Bonferroni’s multiple comparisons test. Period effect *p=0*.*0019*; age effect *p=0*.*1367*; interaction period x age *p=0*.*2912*. For age P7: Baseline *vs* Stim *p=0*.*0436*. For age P8: Baseline *vs* Stim *p=0*.0088 (P6: N=3; P7: N=4; P8: N=5 WT). **h**, same as in **g** with n number of trials. LMM-ANOVA followed by Bonferroni-adjusted post-hoc tests. Period effect *p<0*.*0001*; age effect *p=0*.*1867*; interaction period x age *p=0*.*0122*. For age P6, Stim *vs* Baseline *p<0*.*0001* and Post-stim *vs* Baseline *p=0*.*0018*. For age P7, Stim *vs* Baseline and Stim *vs* Post-stim *have p<0*.*0001*. For age P8; Stim *vs* Baseline *p=0*.*0026* (P6: n=24; P7: n=32; P8: n=40 trials). **i**, Average cool response rate for animals sorted by age. Kruskal Wallis test: *p=0*.*2563* (P6: N=3; P7: N=4; P8: N=5 WT). **j**, same as in **i** with n number of trials. Kruskal Wallis test: *p=0*.*0998* (P6: n=24; P7: n=32; P8: n=40 trials). **k** to **o** follows the same organization as **f** to **j** for the study of the sex effect on GCaMP6f signal of mouse pups receiving thermal cool stimulation. **k**, Top: Peristimulus curve of GCaMP6f signal during thermal cool stimulation (blue box) for animals sorted by sex: Female (F) (light turquoise) and Male (M) (dark turquoise). Bottom: Temperature variations during the cool stimulation. **l**, AUC.s^-1^ quantification for each period and sex. Two-way RM ANOVA followed by Bonferroni’s multiple comparisons test. Period effect *p=0*.*0029*; sex effect *p=0*.*7927*; interaction period x age *p=0*.*9965*. For males: Baseline *vs* Stim *p=0*.*0359*. For females: Baseline *vs* Stim *p=0*.*0059* (females: N=7; males: N=5 WT). **m**, same as in **l** with n number of trials. LMM-ANOVA followed by Bonferroni-adjusted post-hoc tests. Period effect *p< 0*.*0001*; sex effect *p=0*.*7684*; interaction period x sex *p=0*.*7044*. For males, Stim *vs* Baseline *p<0*.*0001* and Stim *vs* Post-stim *p=0*.*0011*. For females, Stim *vs* Baseline *p<0*.*0001*; Stim *vs* Post-stim *p=0*.*007* and Baseline *vs* Post-stim *p=0*.*0161* (females: n=56; males: n=40 trials). **n**, Average cool response rate for animals sorted by sex. Mann-Whitney test *p=0*.*0730* (females: N=7; males: N=5 WT). **o**, same as in **n** with n number of trials. Mann-Whitney test *p=0*.*1654* (females: n=56; males: n=40 trials). Data represented as mean ± sem. Significativity : * p < 0.05; ** p < 0.01; *** p < 0.001; n.s Non-significant.

**Extended Data Fig. 4:**
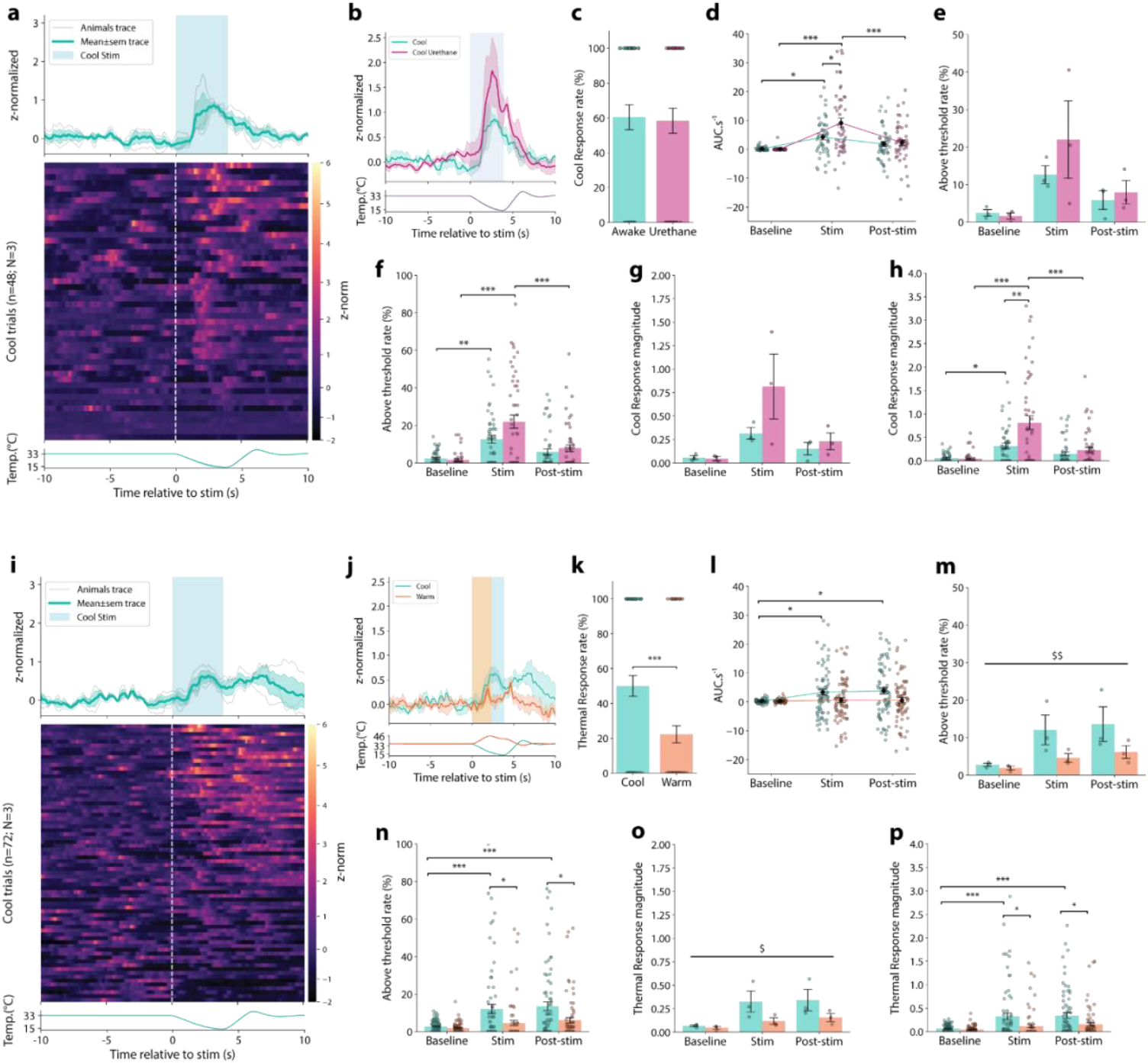
Thermal cool evoked activity in S1. **a**, Top: Peristimulus curve of the mean ± sem GCaMP6f signal during thermal cool stimulation (blue box) for all (blue) and each (grey) animal. Middle: Heatmap of GCaMP6f signal of all cool stimulation (trials) from all animals (n=48 trials in N=3 animals), aligned with the stimulation beginning (white dashed line). Bottom: Temperature variations during the cool stimulation. **b**, Top: Peristimulus curve of the mean ± sem GCaMP6f signal for animals under awake (blue) or urethane (magenta) condition during the thermal cool (blue box). Bottom: Temperature variations during the cool stimulation. **c**, Average cool response rate of awake (blue) or urethane (magenta) animals. Wilcoxon test *p=0*.*8273* (n=48 trials for both conditions). **d**, AUC.s^-1^ quantification for each period of thermal cool stimulation and for the two conditions: awake (blue) and urethane (magenta). LMM-ANOVA followed by Bonferroni-adjusted post-hoc tests. Period effect *p<0*.*0001*; condition effect *p=0*.*156*; interaction period x condition *p=0*.*0144*. For awake condition: Stim *vs* Baseline *p=0*.*0188*. For urethane condition: Stim *vs* Baseline and Stim *vs* Post-stim have *p<0*.*0001*. For the period of stimulation, awake *vs* urethane *p=0*.0264 (n=48 trials for each condition). **e**, Average above threshold rate of the GCaMP6f signal for each period and condition: awake (blue) and urethane (magenta). Two-way RM ANOVA followed by Bonferroni’s multiple comparisons test. Period effect *p=0*.*1809*; condition effect *p=0*.*3238*; interaction period x condition *p=0*.*4851* (N=3 animals for each condition). **f**, same as in **d** with n number of trials. LMM-ANOVA followed by Bonferroni-adjusted post-hoc tests. Period effect *p<0*.*0001*; condition effect *p=0*.*5219*; interaction period x condition *p=0*.*0201*. For awake condition: Stim *vs* Baseline *p=0*.*0012*. For urethane condition: Stim *vs* Baseline and Stim *vs* Post-stim have *p<0*.*0001* (n=48 trials for each condition). **g**, Average cool response magnitude of the GCaMP6f signal for each period and condition: awake (blue) and urethane (magenta). Two-way RM ANOVA followed by Bonferroni’s multiple comparisons test. Period effect *p=0*.*1348*; condition effect *p=0*.*3049*; interaction period x condition *p=0*.*2189* (N=3 WT for each condition). **h**, same as in **f** with n number of trials. LMM-ANOVA followed by Bonferroni-adjusted post-hoc tests. Period effect *p<0*.*0001*; condition effect *p=0*.*1712*; interaction period x condition *p<0*.*0001*. For awake condition: Stim *vs* Baseline *p=0*.*0284*. For urethane condition: Stim *vs* Baseline and Stim *vs* Post-stim have *p<0*.*0001*. For the stimulation: awake *vs* urethane *p=0*.0068 (n=48 trials for each condition). **i** to **p** follow the same organization as **a** to **h** for the comparison of the GCaMP6f signal from mouse pups receiving thermal cool and thermal warm. **i**, Top: Peristimulus curve of the GCaMP6f signal during thermal cool stimulation (blue box) for all (blue) and each (grey) animal. Middle: Heatmap of GCaMP6f signal for all cool stimulation (trials) from all animals (n=48 trials in N=3 WT), aligned with the stimulation beginning (white dashed line). Bottom: Temperature variations during the thermal cool stimulation. **j**, Top: Peristimulus curve of the mean ± sem GCaMP6f signal evoked by thermal cool (blue) or warm (orange) stimulation on the same animals. Bottom: representation of the temperature variations during thermal cool (blue) and warm (orange) stimulation. **k**, Average thermal response rate for cool (blue) and warm (orange) stimulation. Wilcoxon test *p=0*.*0003* (n=72 trials for each condition). **l**, AUC.s^-1^ quantification for each period and condition: cool (blue) and warm (orange). LMM-ANOVA followed by Bonferroni-adjusted post-hoc tests. Period effect *p=0*.*0270;* condition effect *p=0*.*0551*; interaction period x condition *p=0*.*0746*. For cool condition: Stim *vs* Baseline *p=0*.*0119* and Post-stim *vs* Baseline *p=0*.*0442* (n=72 trials for each condition). **m**, Average above threshold rate of the GCaMP6f signal for each period and condition. Two-way RM ANOVA followed by Bonferroni’s multiple comparisons test. Period effect *p=0*.*2562*; condition effect *p=0*.*0098*; interaction period x condition *p=0*.*3199*. The main effect of the condition is noted as $$ (N=3 WT for each condition). **n**, same as in **m** with n number of trials. LMM-ANOVA followed by Bonferroni-adjusted post-hoc tests. Period effect *p<0*.*0001*; condition effect *p=0*.*0133*; interaction period x condition *p=0*.*0707*. For cool condition: Stim *vs* Baseline and Post-stim *vs* Baseline have *p<0*.*0001*. For the stimulation period: cool *vs* warm *p=0*.*0246*. For the Post-stim period: cool *vs* warm *p=0*.*0251* (n=72 trials for each condition). **o**, Average thermal response magnitude of the GCaMP6f signal for each period and condition. Two-way RM ANOVA followed by Bonferroni’s multiple comparisons test. Period effect *p=0*.*2609*; condition effect *p=0*.*0118*; interaction period x condition *p=0*.*3039*. The main effect of the condition is noted as $ (N=3 WT for each condition). **p**, same as in **o** with n number of trials. LMM-ANOVA followed by Bonferroni-adjusted post-hoc tests. Period effect *p<0*.*0001*; condition effect *p=0*.*0041*; interaction period x condition *p=0*.*0393*. For cool condition: Stim *vs* Baseline and Post-stim *vs* Baseline have p<0.0001. For the stimulation period: cool *vs* warm *p=0*.*0073*. For the Post-stim period: cool *vs* warm *p=0*.0221 (n=72 trials for each condition). Data represented as mean ± sem. Significativity : * p < 0.05; ** p < 0.01; *** p < 0.001; n.s Non-significant.

**Extended Data Fig. 5:**
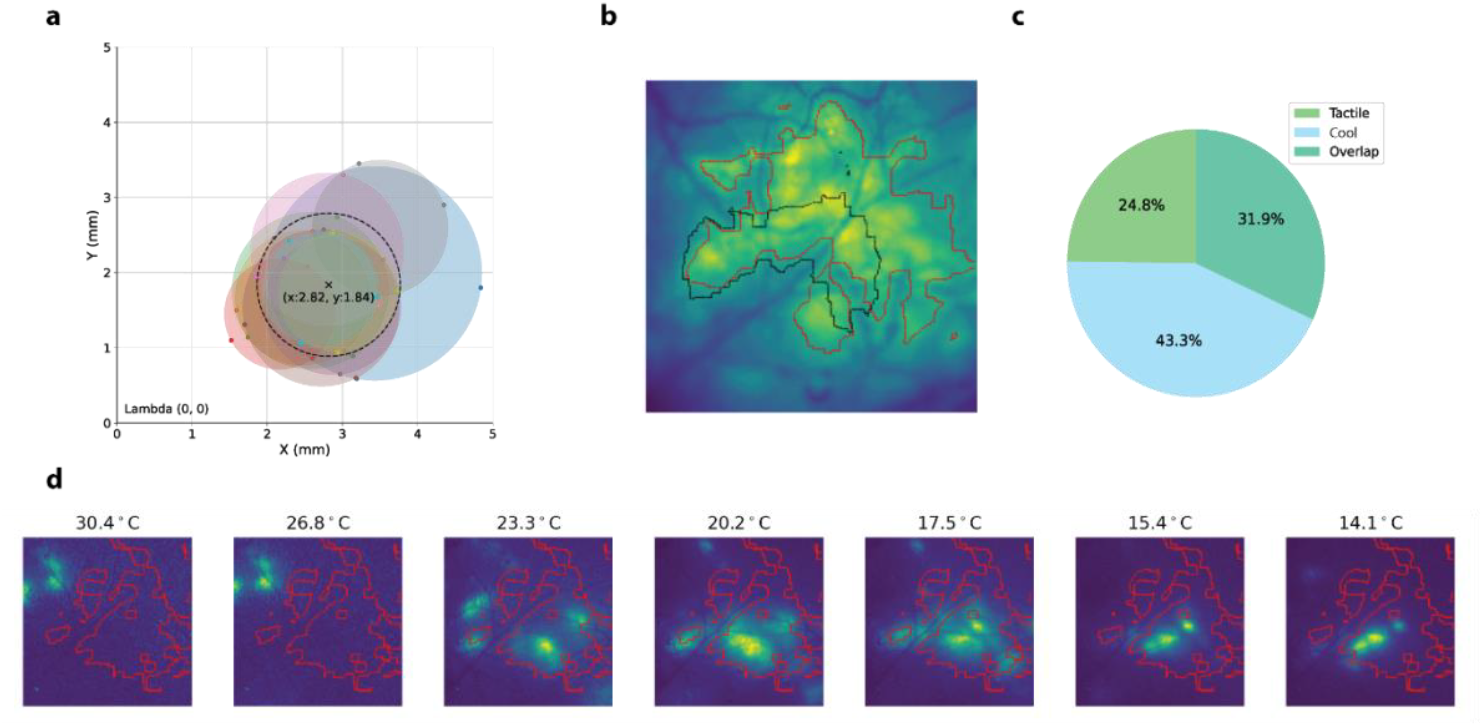
Overlap of cool and tactile cortical receptive fields. **a**, Mapping of stereotaxic coordinates from animal recording areas with the mean x and y coordinates from the lambda. **b**, Example of tactile (black) and thermal cool (red) receptive field. **c**, Quantification of the cumulative tactile and cool receptive field proportion of the tactile and cool specific receptive field and their overlapping. **d**, Example of cortical activation kinetics for a cool stimulation.

**Extended Data Fig. 6:**
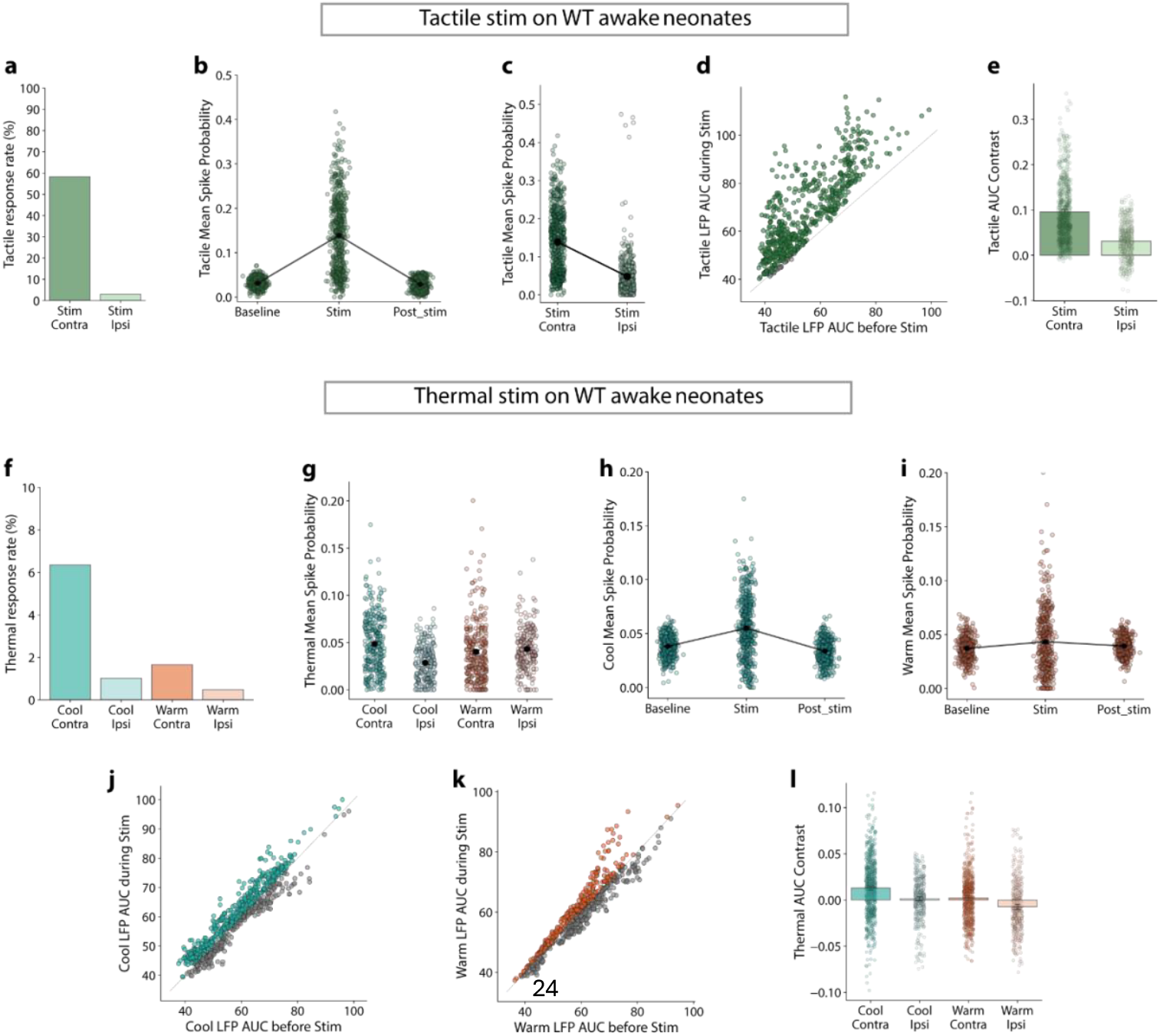
WT cortical tactile and thermal evoked activity. **a**, Contingency analysis of the tactile response rate for contra- and ipsilateral stimulation (n_contra_ = 566; n_ipsi_ = 401 clusters). **b**, Mean tactile spike probability quantification for the three recording periods. (n=566 clusters). **c**, Mean tactile spike probability quantification for the contra- and the ipsilateral stimulation. (n_contra_=566 and n_ipsi_=401 clusters). **d**, Quantification of the tactile AUC of the derived LFP signal for the Baseline and Stim periods. Each point represents a channel. (n=608 channels).**e**, Quantification of the tactile AUC-derived LFP signal contrast (Stim to Baseline signal) for the contra- or ipsilateral side. Each point represents a channel. (n_contra_=608 and n_ipsi_=480 channels). **f**, Contingency analysis of the thermal response rate for cool, warm, contralateral, and ipsilateral stimulations. (n_cool-contra_=472, n_cool-ipsi_=293, with n_warm-contra_=415, n_warm-ipsi_=209 clusters). **g**, Mean thermal spike probability quantification for cool, warm, contralateral, and ipsilateral stimulations (n_cool-contra_=289, n_cool-ipsi_=287, with n_warm-contra_=204, n_warm-ipsi_=210 clusters). **h**, Mean cool spike probability quantification for the three periods of recording (n=504 clusters). **i**, Same as **h** for warm stimulations (n=422 clusters). **j**, Cool AUC-derived LFP signal quantification for the Baseline and Stim periods. Each point represents a channel. (n=704 channels). **k**, same as **j** for warm stimulation. (n=608 channels). **l**, Quantification of the thermal AUC-derived LFP signal contrast (Stim to Baseline) for the cool, warm contra- and ipsilateral stimulation. Each point represents channel (n_cool-contra_=704, n_cool-ipsi_=416, with n_warm-contra_=608, n_warm-ipsi_=320 channels). Data represented as mean ± sem.

**Extended Data Fig. 7:**
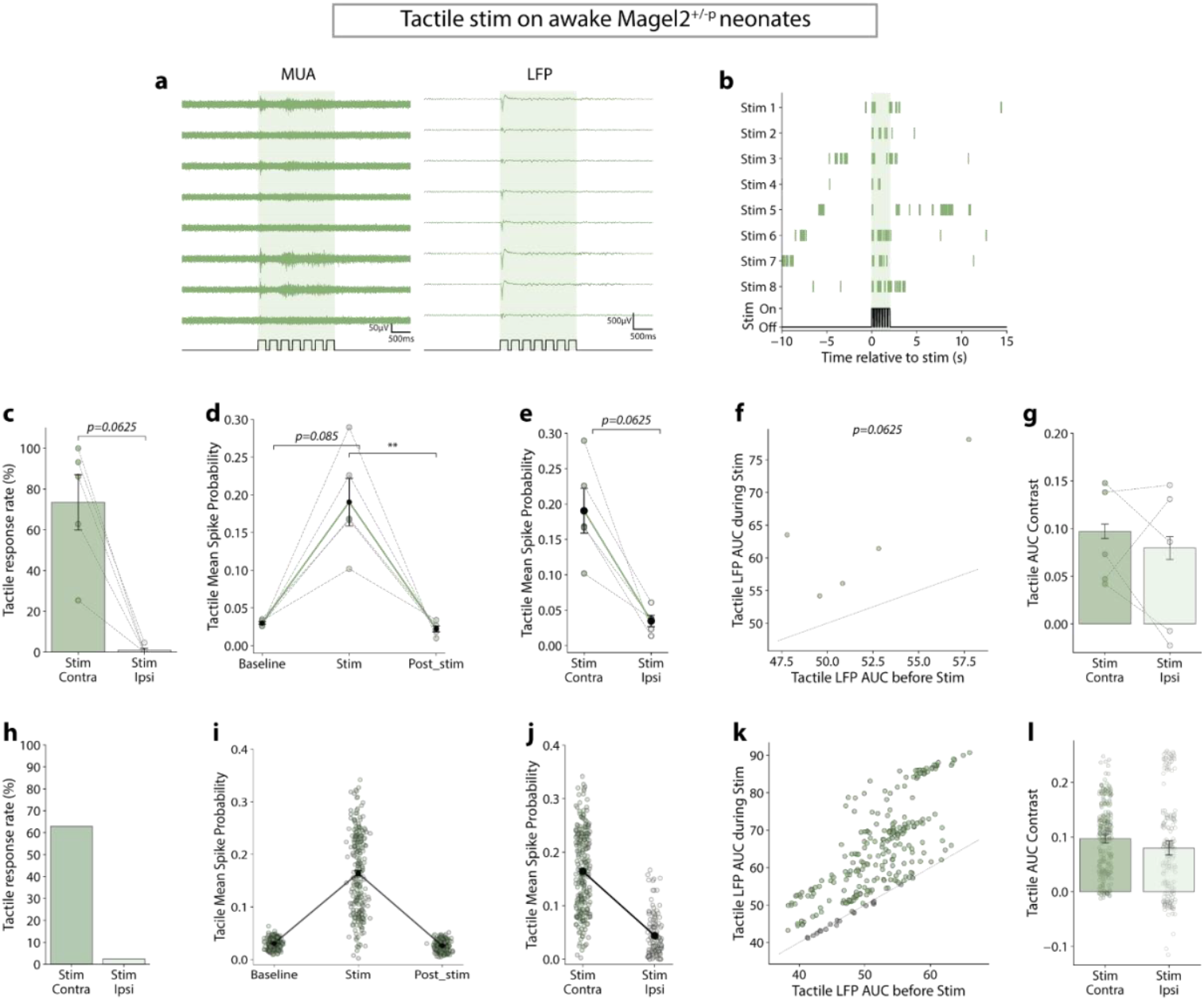
S1 tactile activity coding in Magel2 KO newborns. **a**, Representative tactile evoked MUA and LFP for a tactile stimulation (green box) in the S1 of an awake Magel2 KO newborn. Traces from one shank are organized from the shallowest to the deepest channel. **b**, Raster plot of an example cluster with the 8 tactile stimuli (green box). **c**, Quantification of the mean tactile response rate for the contra- and the ipsilateral side. Wilcoxon test *p=0*.*0625* (N=5 Magel2 KO for both conditions). **d**, Mean tactile spike probability quantification for the three periods of recording. Friedman test *p=0*.*0150* followed by Dunn post-hoc with Bonferroni adjustment: Baseline *vs* Stim *p=0*.*0851*, Stim *vs* Post-stim *p=0*.*0056* (N=5 Magel2 KO). **e**, Mean tactile spike probability quantification for the contra- and the ipsilateral side. Wilcoxon test *p=0*.*0625* (N = 5 Magel2 KO for both conditions). **f**, Quantification of the tactile AUC of the derived LFP signal for the Baseline and Stim periods. Each point represents an animal. Wilcoxon test *p=0*.*0625* (N=5 Magel2 KO). **g**, Quantification of the contrast of the tactile AUC-derived LFP signal (Stim to Baseline) for the contra- or ipsilateral side. Each point represents one channel. Wilcoxon test *p=0*.*0015* (N=5 Magel2 KO). **h** to **i** followed the same organization as **c** to **g** with a number of clusters or channels instead of animals. **h**, (n_contra_=327, n_ipsi_=123 clusters). **i**, (n=237 clusters). **j**, (n_contra_=327, n_ipsi_=123 clusters). **k**, (n=288 channels). **l**, (n_contra_=288, n_ipsi_=192 channels). Data represented as mean ± sem. Significativity : * p < 0.05; ** p < 0.01; *** p < 0.001; n.s Non-significant.

**Extended Data Fig. 8:**
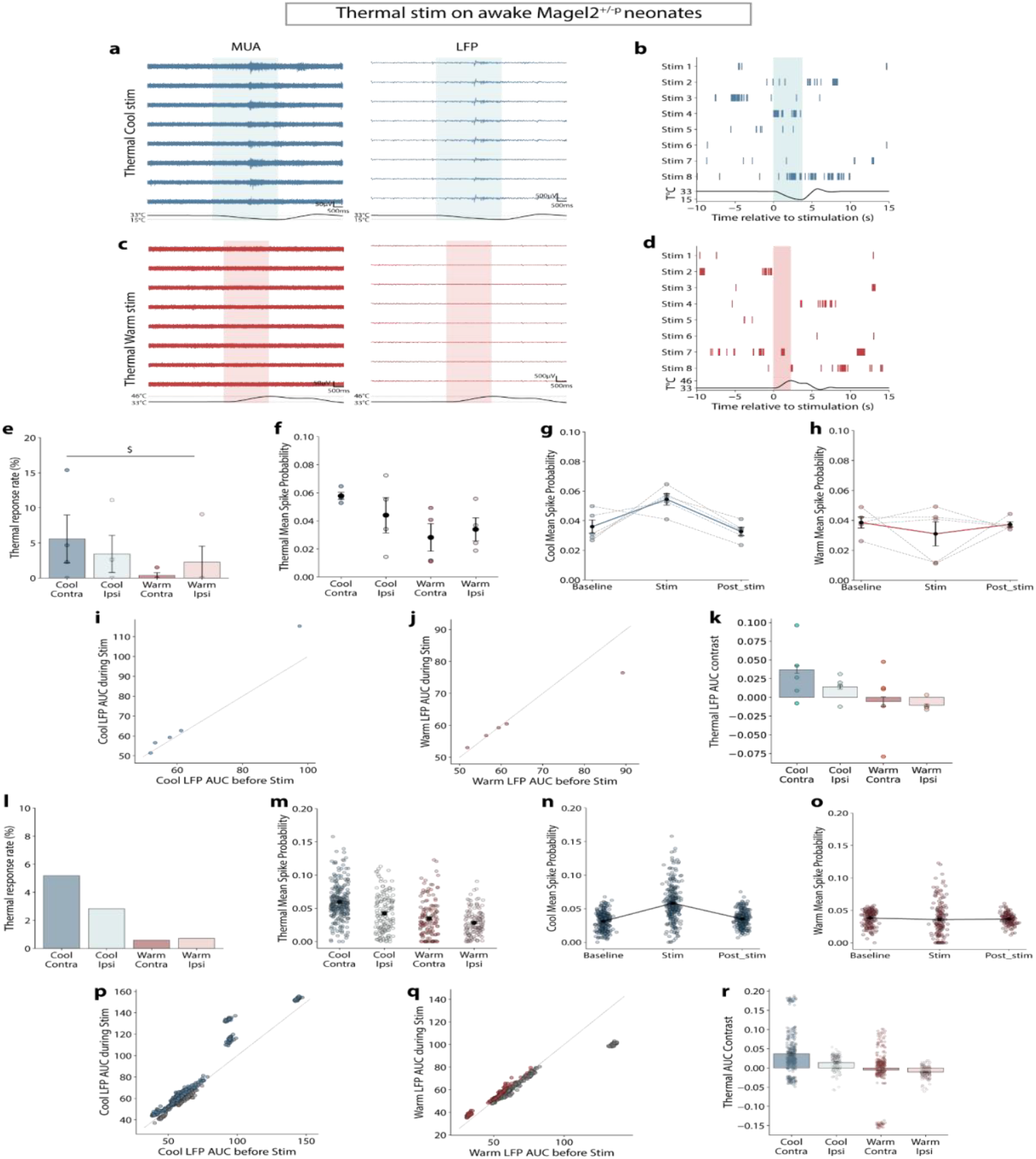
S1 thermal activity coding in Magel2 KO newborns. **a**, Representative thermal evoked MUA and LFP for a cool stimulation (blue box) in the S1 cortex of an awake Magel2 KO newborn. Traces from one shank are organized from the shallowest to the deepest channel. **b**, Raster plot of an example cluster with the 8 thermal cool stimuli (blue box). **c**, same as **a** for the warm stimulation (red box). **d**, same as **b** for warm stimulation (red box). **e**, Mean thermal response rate quantification for cool, warm, contra-, and ipsilateral stimulations. Two-way RM ANOVA followed by Bonferroni’s multiple comparisons test. Temperature effect *p=0*.*0328*; laterality effect *p=0*.*9533*; interaction temperature x laterality *p=0*.*6295*. Any comparisons were significant for Bonferroni’s multiple comparisons test. Main effect of the temperature noted as $. (N=4 Magel2 KO for all conditions). **f**, Mean thermal spike probability quantification for cool, warm, contralateral, and ipsilateral stimulations. Two-way RM ANOVA followed by Bonferroni’s multiple comparisons test. Temperature effect *p=0*.*1355*; laterality effect *p=0*.*5986*; interaction temperature x laterality *p=0. 4488*. (N=4 Magel2 KO for all conditions). **g**, Mean cool spike probability quantification for the three periods of recording. Friedman test *p=0*.*0907* followed by Dunn post-hoc with Bonferroni adjustment: (N=5 Magel2 KO). **h**, Mean warm spike probability quantification for the three periods of recording. Friedman test *p=0*.*5488*. (N=5 Magel2 KO). **i**, Cool AUC of the derived LFP signal quantification for the Baseline and Stim periods. Each point represents an animal. Wilcoxon test *p=0*.*125* (N=5 Magel2 KO). **j**, same as **i** for warm stimulation. Wilcoxon test *p=0*.*8125* (N=5 Magel2 KO). **k**, Quantification of the thermal AUC-derived LFP signal contrast (Stim to Baseline) for the cool, warm contra- and ipsilateral stimulation. Each point represents an animal. Mixed-effects analysis followed by Bonferroni’s multiple comparisons test. Temperature effect *p=0*.*1518*; laterality effect *p=0*.*3707*; interaction temperature x laterality *p=0*.*6123*. (N=5 Magel2 KO). **l** to **r** follow the same organization as **e** to **k** with a number of cluster or channel instead of animals. **l**, (n_cool-contra_=274, n_cool-ipsi_=137, with n_warm-contra_=169, n_warm-ipsi_=137 clusters). **m**, (n_cool-contra_=274, n_cool-ipsi_=143, with n_warm-contra_=141, n_warm-ipsi_=137 clusters). **n**, (n=289 clusters). **o**, (n=170 clusters). **p**, (n=416 channels). **q**, (n=256 channels). **r**, (n_cool-contra_=416, n_cool-ipsi_=160, with n_warm-contra_=256, n_warm-ipsi_=192 channels). Data represented as mean ± sem. Significativity : * p < 0.05; ** p < 0.01; *** p < 0.001; n.s Non-significant.

**Extended Data Fig. 9:**
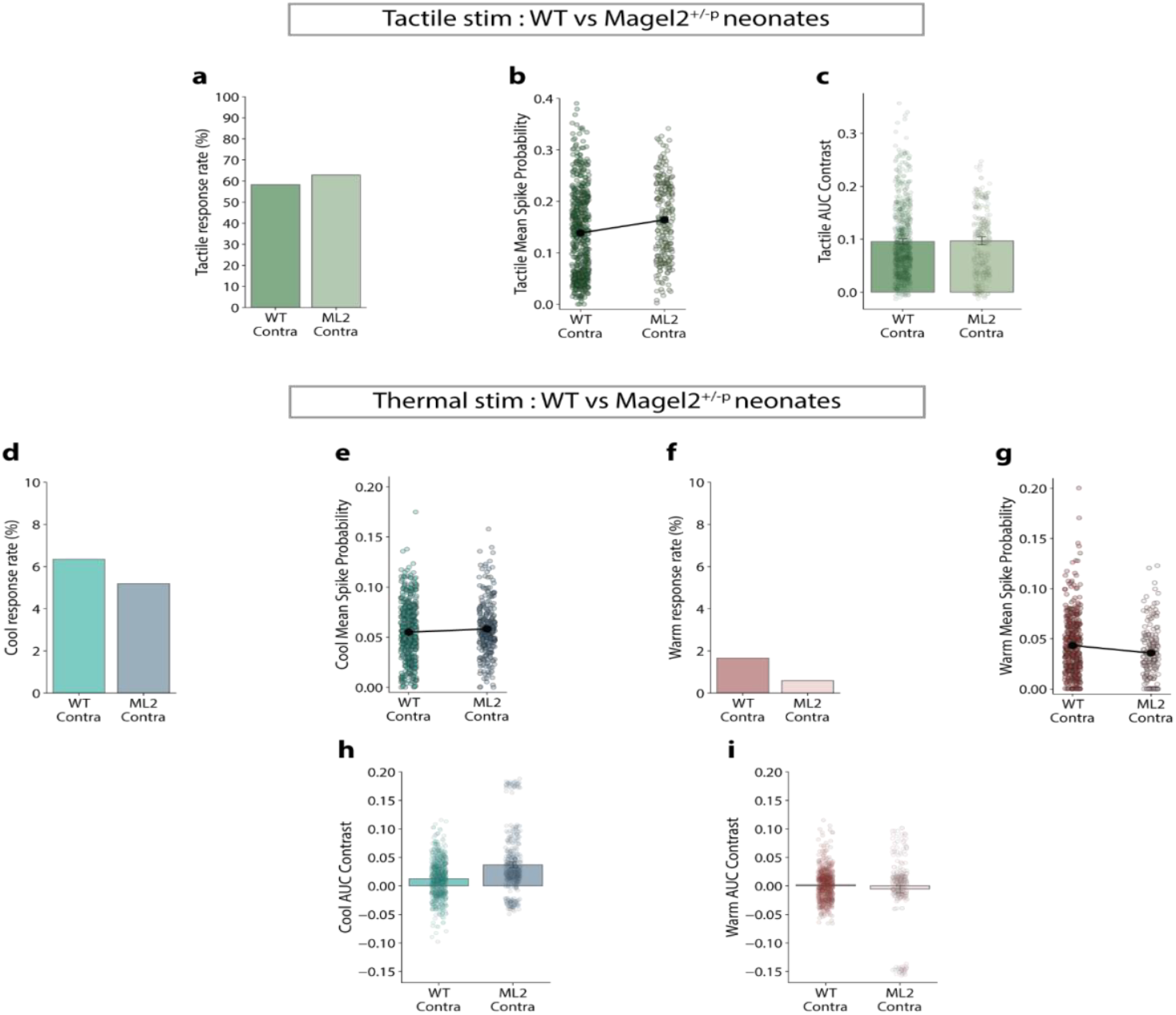
WT vs Magel2 KO neonates somatosensory evoked activity. **a**, Contingency analysis of the tactile response rate quantification for WT and Magel2 KO neonates (n_WT_ =566 and n_Magel2_=237 clusters). **b**, Mean tactile spike probability quantification for WT and Magel2 KO neonates (n_WT_ =566 and n_Magel2_=237 clusters). **c**, Quantification of the tactile AUC-derived LFP signal contrast (Stim to Baseline) for WT and Magel2 KO neonates. Each point represents one channel (n_WT_ =608 and n_Magel2_=288 channels). **d**, Mean thermal cool response rate for WT and Magel2 KO neonates (n_WT_ =504 and n_Magel2_=289 clusters). **e**, Mean thermal cool spike probability for WT and Magel2 KO neonates (n_WT_ =504 and n_Magel2_=289 clusters). **f**, Same as **d** for warm stimulation (n_WT_ =422 and n_Magel2_=170 clusters). **g**, Same as **e** for warm stimulation (n_WT_ =422 and n_Magel2_=170 clusters). **h**, Quantification of the cool AUC-derived LFP signal contrast for WT and Magel2 KO neonates. Each point represents one channel (n_WT_ =704 and n_Magel2_=416 channels). **i**, same as **h** for warm stimulation (n_WT_ =608 and n_Magel2_=256 channels). Data represented as mean ± sem.

